# A Global Ligandability Map of Tryptoline Butynamide Stereoprobes Identifies Covalent Inhibitors of the Actin Maturation Protease ACTMAP

**DOI:** 10.64898/2026.02.21.707170

**Authors:** Yijun Xiong, Christopher J. Reinhardt, Tracey Nguyen, Melissa A. Hoffman, Gabriel M. Simon, Bruno Melillo, Benjamin F. Cravatt

**Affiliations:** Department of Chemistry, The Scripps Research Institute, La Jolla, California 92037, United States; Vividion Therapeutics, San Diego, California 92121, United States

## Abstract

Covalent chemistry coupled with activity-based protein profiling (ABPP) offers a versatile approach for small-molecule ligand discovery in native biological contexts. The covalent ligandability maps generated by ABPP that target cysteine have frequently leveraged the acrylamide as a reactive group due to its tempered electrophilicity and presence in many advanced tool compounds and therapeutics. More recently, alternative cysteine-directed reactive groups such as the butynamide have emerged as an additional source of covalent probes and drugs, but their global reactivity with the proteome remains largely unexplored. Here, we compare the ligandability maps of stereochemically defined acrylamide and butynamide compounds (stereoprobes) built from a common tryptoline core and find that the butynamides, despite exhibiting attenuated intrinsic and proteome-wide reactivity, preferentially engage a diverse set of proteins in human cancer cells. Among the butynamide-preferring proteins was C19orf54/ACTMAP, a cysteine protease required for the post-translational maturation of actin. We show that (1*S*, 3*R*)-tryptoline butynamides stereoselectively react with the catalytic nucleophile of ACTMAP, leading to accumulation of *N*-terminally unprocessed actin in cancer cells. Our findings support reactive group diversification as a strategy for expanding the ligandability of the human proteome and the butynamide, more specifically, as a differentiated cysteine-directed electrophile for chemical probe discovery.

---

Small molecules are valuable tools to perturb the functions of proteins in biological systems and a major category of therapeutics. The systematic pursuit of chemical probes, however, must grapple with the structural diversity of human proteins, many of which also lack robust functional assays for small-molecule screening.^1, 2^ To address such challenges, innovative approaches like fragment-based drug discovery,^3, 4^ DNA-encoded libraries,^5, 6^ and activity-based protein profiling (ABPP)^7–10^ have emerged to explore small molecule–protein interactions across a wide range of proteins.^11^ Among these strategies, the chemical proteomic method ABPP has an advantage of evaluating the small molecule binding potential (or ligandability) of proteins in native biological systems and thus can account for the myriad post-transcriptional and post-translational mechanisms that regulate protein structure and function.

In addition to the advent of generalizable assays for measuring small molecule-protein interactions, the types of chemistry screened by such assays offers another source of innovation for expanding the ligandable proteome. Features of small-molecule library design that can influence protein binding include compound size (fragments vs macrocycles),^12–14^ structure (sp^2^ vs sp^3^-rich scaffolds),^15^ and nature of bonding (non-covalent vs covalent).^16^ Over the past two decades, covalent chemistry has emerged as a particularly rich source of chemical probes and drugs and can provide a way to target proteins with greater selectivity (e.g., by targeting isoform-restricted nucleophilic residues)^17–19^ and durability (e.g., by coupling the length of pharmacodynamic effects to protein turnover),^20, 21^ as well as to identify ligands for cryptic and dynamic pockets on proteins.^9^ The integration of covalent chemistry with ABPP has further enabled global ligand discovery in living cells, leading to the identification of chemical tools that target a diverse array of proteins, including enzymes,^7, 22–28^ RNA-binding proteins,^29–32^ transcription factors^33–36^, E3 ligase systems,^37–40^ and adaptors.^41, 42^ In several instances, the ligands have been found to engage proteins by nonconventional mechanisms (e.g., ATP-cooperative,^25^ DNA-dependent,^33^ or complexoform-restricted^43^ interactions) that underscore the range of factors that can influence the small molecule-binding potential of proteins in cells.

While we and others have been interested in evaluating covalent chemistry approaches that target different nucleophilic amino acids,^44–61^ the most common category of advanced covalent chemical probes and drugs acts by targeting cysteine.^10, 20, 62–66^ Among cysteine-directed electrophiles, the acrylamide is often favored,^62, 67^ likely due to its tempered reactivity and small size, which has facilitated its use in converting reversible inhibitors into irreversible probes and drugs, as well as in library designs for covalent-first ligand discovery.^13, 42, 62, 68, 69^ Our lab has specifically leveraged the acrylamide in the construction of diversity-oriented synthesis-inspired sets of stereochemically defined electrophilic compounds (or ‘stereoprobes’), which we have deployed in ABPP studies to generate covalent ligandability maps of primary human immune cells^40, 70^ and human cancer cell lines.^34, 41–43^ These experiments have identified hundreds of stereoselective liganding events across the human proteome, supporting the versatility of the acrylamide for covalent chemical probe discovery. Nonetheless, the extent to which stereoprobes bearing alternative cysteine-directed electrophiles may engage overlapping or distinct proteins remains largely unexplored. Here, we set out to address this question by generating and characterizing a set of tryptoline stereoprobes where the acrylamide has been replaced with a structurally related butynamide, which is also found in specific drugs and drug candidates.^17, 71, 72^ Leveraging an ABPP protocol for quantitative comparison of tryptoline butynamide and acrylamide stereoprobe reactivity in pooled sets of cancer cell lines, we identified several proteins that were preferentially or exclusively engaged by butynamides. We confirmed the butynamide-preferred liganding of multiple proteins through recombinant expression and characterization, including the cytosolic cysteine protease ACTMAP, which is responsible for cleaving the *N*-acetylated methionine of actin.^73^ We show that tryptoline butynamides react with the conserved cysteine nucleophile in ACTMAP, leading to stereoselective blockade of actin maturation in cells. Our findings highlight the importance of electrophile identity in shaping the global and specific reactivity of covalent small molecules, as well as the potential to expand the ligandability of the proteome by leveraging cysteine-directed reactive groups beyond the acrylamide.

## Results and Discussion

### Generation and initial characterization of tryptoline butynamide stereoprobes

Previous ABPP studies have shown that tryptoline acrylamide stereoprobes engage a large and diverse array of proteins in human cells.^42, 70^ We therefore selected the tryptoline core for constructing a matching set of butynamide stereoprobes, where both the tryptoline acrylamides and butynamides were generated in parent (non-alkyne) and alkyne-modified form (compounds **1-16**; **Figure 1A**). We initially compared the global proteomic reactivity of the alkyne-modified tryptoline acrylamides (WX-01-05/06/07/08 (**5-8**)) and butynamides (WX-02-568/570/569/571 (**13-16**)) (10 µM, 1 h) in the Ramos B-lymphocyte human cancer cell line by gel-ABPP methods.^42^ The tryptoline butynamides showed lower overall proteomic reactivity compared to the tryptoline acrylamides (**Figures 1B** and **S1**), which also matched the respective glutathione reactivities of the two sets of stereoprobes (**Table S1**). These results are consistent with previous work showing that butynamides generally display attenuated electrophilicity compared to structurally related acrylamides.^74, 75^

**Figure 1.**
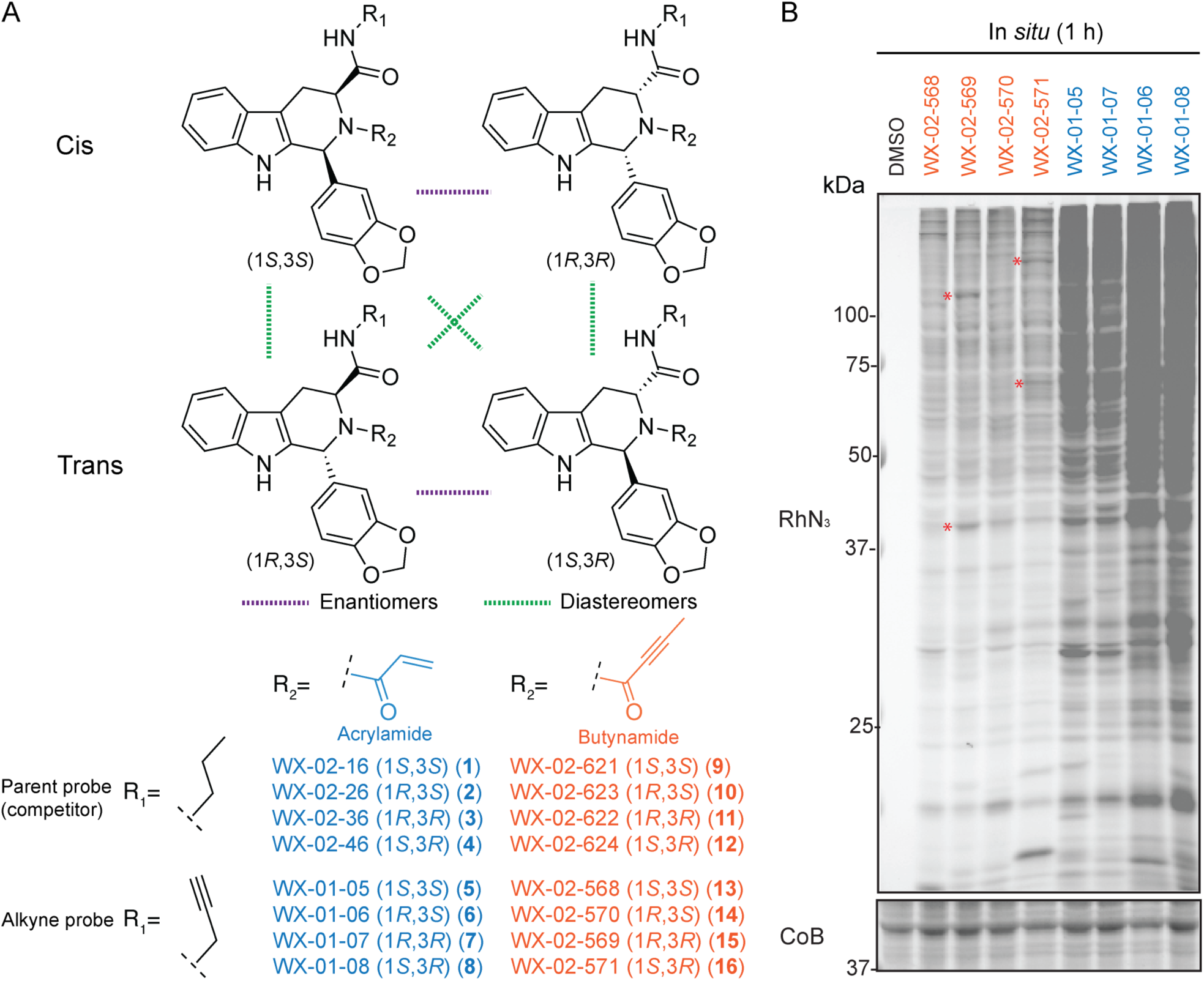
Design and initial profiling of tryptoline acrylamide and butynamide stereoprobes. (A) Structures of tryptoline acrylamide^42^ and butynamide stereoprobes. (B) Gel-ABPP data for Ramos cells treated with the indicated alkyne stereoprobes (10 µM) for 1 h, followed by lysis and visualization of stereoprobe-reactive proteins by copper-catalyzed azide-alkyne cycloaddition (CuAAC) conjugation^76, 77^ to an azide-rhodamine (RhN3) reporter group, SDS-PAGE, and in gel fluorescence scanning.^105^ CoB, Coomassie blue staining (shorter exposure time for gel-ABPP data provided in **Figure S1**). The names of tryptoline butynamides and acrylamides are shown in orange and blue font, respectively. Red asterisks mark representative proteins that were stereoselectively engaged by tryptoline butynamides. Data are from a single experiment representative of two independent experiments.

### Mapping proteins liganded by tryptoline butynamide stereoprobes

The gel-ABPP experiments also detected several stereoselective tryptoline butynamide-protein interactions in Ramos cells (**Figure 1B**, red asterisks). We next set out to identify proteins that were stereoselectively liganded by the tryptoline butynamides and acrylamides using protein-directed ABPP,^42^ which involves treating cells with parent stereoprobe competitors (acrylamides: WX-02-16/26/36/46 (**1-4**); butynamides: WX-02-621/622/623/624 (**9-12**) (20 µM, 2 h)) or DMSO control, followed by stereochemically matched alkynes (acrylamides: WX-01-05/06/07/08 (**5-8**); butynamides: WX-02-568/569/570/571 (**13-16**) (5 µM, 1 h)) (**Figure S2A**). Cells are then lysed and alkyne stereoprobe-modified proteins conjugated to biotin-azide by copper-catalyzed azide-alkyne cycloaddition (CuAAC) chemistry,^76, 77^ enriched with streptavidin beads, digested with trypsin on beads, and analyzed by multiplexed (tandem mass tag (TMT)) mass spectrometry (MS)-based proteomics. Protein-directed ABPP has previously been applied to individual cell lines,^41, 42^ but here we adapted the method to analyze a pooled collection of five human cancer cell lines from distinct lineages (**Figure S2A**) toward the goal of broadening the proteomic coverage of the stereoprobe-protein interaction (or ‘ligandability’) maps.

Proteins were considered liganded by stereoprobes if they showed > three-fold enantioselective enrichment by an alkyne-modified stereoprobe, and this enrichment was blocked > two-fold by the corresponding parent stereoprobe competitor. Both the tryptoline butynamides and acrylamides liganded a diverse array of proteins (**Dataset S1**), and these liganding events were distributed across all four stereo-isomers, as visualized in quadrant plots (**Figures 2A** and **S2B**). The tryptoline acrylamide ligandability maps included proteins previously found to be liganded in protein-directed ABPP experiments performed with individual cells lines.^33, 42^ Some of these proteins also displayed restricted expression across the five cell lines examined herein (**Figure S3**), indicating that the pooled format for protein-directed ABPP maintained a suitable level of sensitivity.

**Figure 2.**
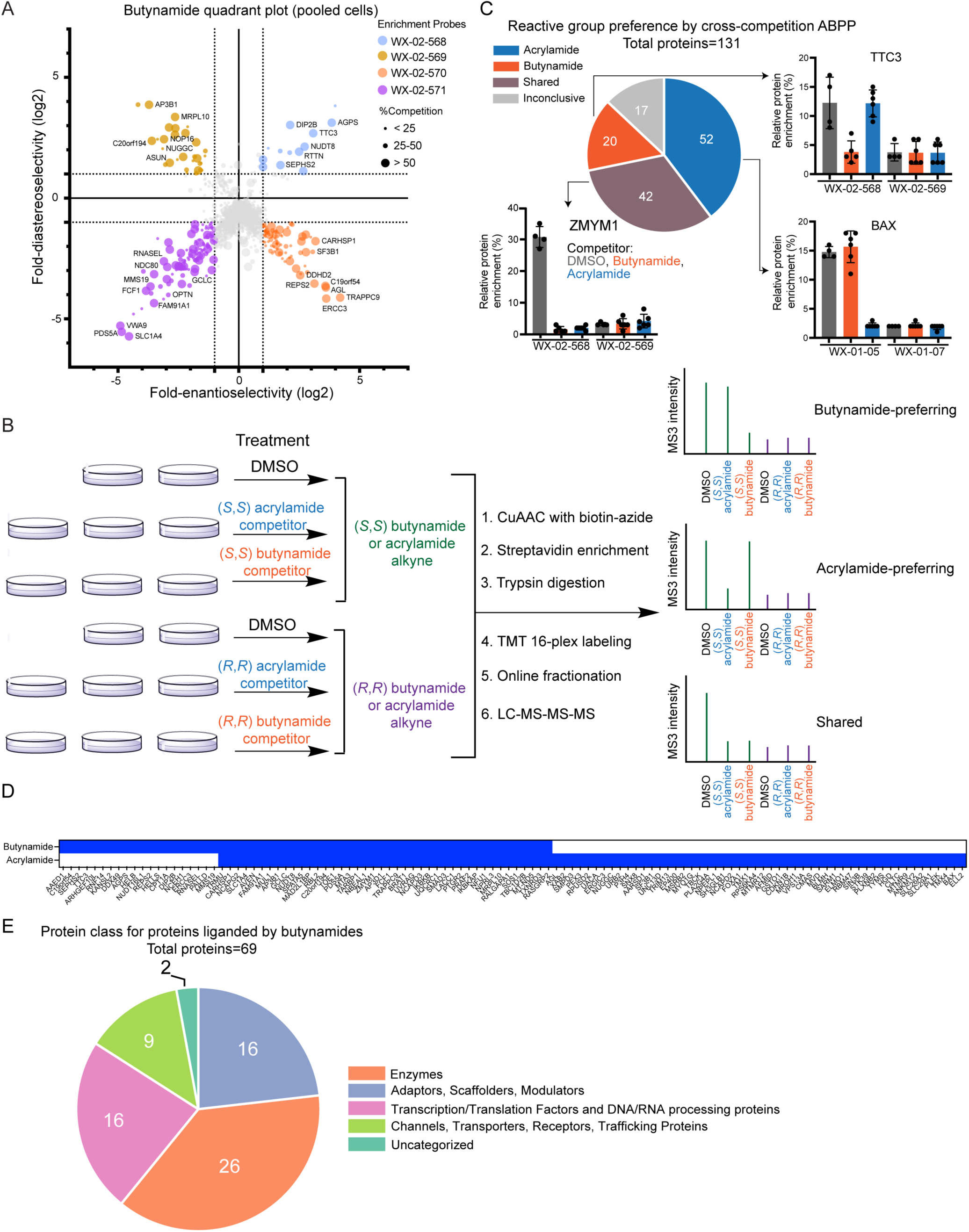
Mapping proteins preferentially liganded by tryptoline butynamides and acrylamides by pro-tein-directed ABPP. (A) Quadrant plot highlighting stereoselectively liganded proteins for each stereocon-figuration of the tryptoline butynamides as determined by protein-directed ABPP in pooled human cancer cell lines. Enantioselectivity (x axis) is the ratio of enrichment for one stereoisomer versus its enantiomer, and diastereoselectivity (y axis) is the ratio of enrichment of one stereoisomer versus the average of its two diastereomers. Pooled cell lines were treated with DMSO or parent tryptoline butynamides (20 µM) for 2 h followed by the stereomatched alkyne tryptoline butynamide (5 µM) for 1 h and analyzed by protein-directed ABPP (also see **Figure S2A**). (B) Workflow of cross-competition protein-directed ABPP, in which pooled cell lines were pretreated with parent tryptoline acrylamides or butynamides (competi-tors; 20 µM) for 2 h followed by the stereomatched alkyne tryptoline acrylamides or butynamides (5 µM) for 1 h. Mock data on the right show examples of butynamide-preferring, acrylamide-preferring, and shared liganded proteins. (C, D) Pie chart (C) and heat map (D) showing distribution of butynamide-preferring, acrylamide-preferring, and shared liganded proteins. For C, examples of acrylamide-preferring (BAX), butynamide-preferring (TTC3), and shared (ZMYM1) proteins are shown. For D, blue color de-notes the proteins in each liganding category (proteins with inconclusive reactive group preferences from the cross-competition ABPP experiments (17 total) were not included in the heat map). See Results section for description of parameters used to define each category of liganding event. (E) Functional class distribution of butynamide-liganded proteins assigned as described previously^42, 70^ using GO (Panther) and Uniprot annotations. For A and D, the protein-directed ABPP data represent average values from four to six independent experiments per stereoprobe. For D, error bars represent s.d. values.

To more directly compare the protein targets of tryptoline butynamides and acrylamides, we adapted the protein-directed ABPP protocol for cross-competition analysis such that the pooled cancer cell lines were first treated with stereo-matched parent tryptoline acrylamides and butynamides (e.g., WX-02-16 or WX-02-621) followed by the corresponding alkyne-modified tryptoline acrylamide or butynamide (e.g., WX-01-05 or WX-02-568), and the samples were then combined for analysis in the same multiplexed MS-based experiment (**Figure 2B**). In this cross-competition format, each stereo-matched parent acrylamide and butynamide is directly compared for their respective blockade of protein enrichment by the corresponding alkyne-modified acrylamide or butynamide, thus accounting for all proteins that were stereoselectively enriched by either reactive group. Liganded proteins were then categorized as follows: 1) butynamide-preferring, if showing > two-fold blockade of enrichment by the parent butynamide that was also > 1.5-fold more than the blockade by the stereo-matched parent acrylamide; 2) acrylamide-preferring, if showing > two-fold blockade of enrichment by the parent acrylamide that was also > 1.5-fold more than the blockade by the stereo-matched parent butynamide; or 3) shared, if showing > two-fold and near-equivalent blockade of enrichment by both the parent butynamide and acrylamide (**Figure 2B**).

More than half of the ∼130 liganded proteins showed a clear preference for engaging acrylamides (52 total proteins) or butynamides (20 total proteins) (**Figure 2C, D**), underscoring the impact of the reactive group on shaping stereoprobe-protein interactions. These results also revealed that the tryptoline butynamides, despite exhibiting lower glutathione (**Table S1**) and overall proteomic (**Figure 1B**) reactivity compared to tryptoline acrylamides, engaged a number of unique proteins (**Figure 2C, D**). The tryptoline butynamides also avoided specific targets, such as the spliceosome factor SF3B1 (**Figure S4A**), for which enantioselective liganding by both (1*R*, 3*S*) and (1*R*, 3*R*) tryptoline acrylamides has been found to produce general anti-proliferative effects.^41^ Accordingly, we did not observe stereoselective proliferation defects in cells treated with (1*R*, 3*S*) and (1*R*, 3*R*) tryptoline butynamides (**Figure S4B**).

The tryptoline butynamide-liganded proteins were distributed across diverse structural and functional classes, including categories that have been historically challenging to target with small molecules (e.g., transcriptional regulatory proteins and adaptor/scaffolding proteins) (**Figure 2E**). We next sought to map the liganded cysteines in proteins engaged by tryptoline butynamides using cysteine-directed ABPP.^70, 78^ These experiments mapped > 12,000 cysteines in the pooled cell samples, but only a modest number of butynamide-liganded cysteines were identified (23 in total; defined as showing > 50% enantioselective decreases in iodoacetamide-desthiobiotin reactivity; **Dataset S1**). We suspect that cysteine-directed ABPP may suffer a greater loss in sensitivity than protein-directed ABPP when performed with pooled cell line samples, as the former method requires detection and quantification of individual (vs aggregate) peptides for each liganded protein. We next sought to experimentally confirm and further characterize a representative set of butynamide-preferring proteins.

### Further characterization of proteins preferentially liganded by tryptoline butynamides

We initially selected two butynamide-preferring proteins – AGPS and HELLS – for further characterization. AGPS, or alkyldihydroxyacetonephosphate synthase, is a flavin adenine dinucleotide (FAD)-dependent enzyme involved in ether lipid metabolism^79–81^ and was found to be preferentially liganded by (1*S*, 3*S*) alkyne/parent tryptoline butynamide pair WX-02-568/WX-02-621 (**Figure 3A**). The cysteine-directed ABPP data assigned C190 as the site of engagement by WX-02-621 with other quantified cysteines in AGPS being unaltered (**Figures 3B**). We confirmed by gel-ABPP that WX-02-568 stereoselectively reacted with recombinant WT-AGPS expressed in HEK293T cells, and this interaction was stereoselectively blocked by pre-treatment with WX-02-621, but not by the stereomatched tryptoline acrylamide WX-02-16 (**Figure 3C**). In contrast, a C190A-AGPS mutant did not react with WX-02-568 (**Figure 3C**). Based on a crystal structure of guinea pig AGPS (PDB: 5ADZ), C190 is located near the FAD-binding site, but distal to the sites of binding of established active-site directed inhibitors of AGPS (**Figure 3D**).^81, 82^ These findings suggest that covalent ligands targeting C190 of AGPS may have the potential to allosterically modulate the activity of this enzyme in a manner that is distinct from established inhibitors.^82^

**Figure 3.**
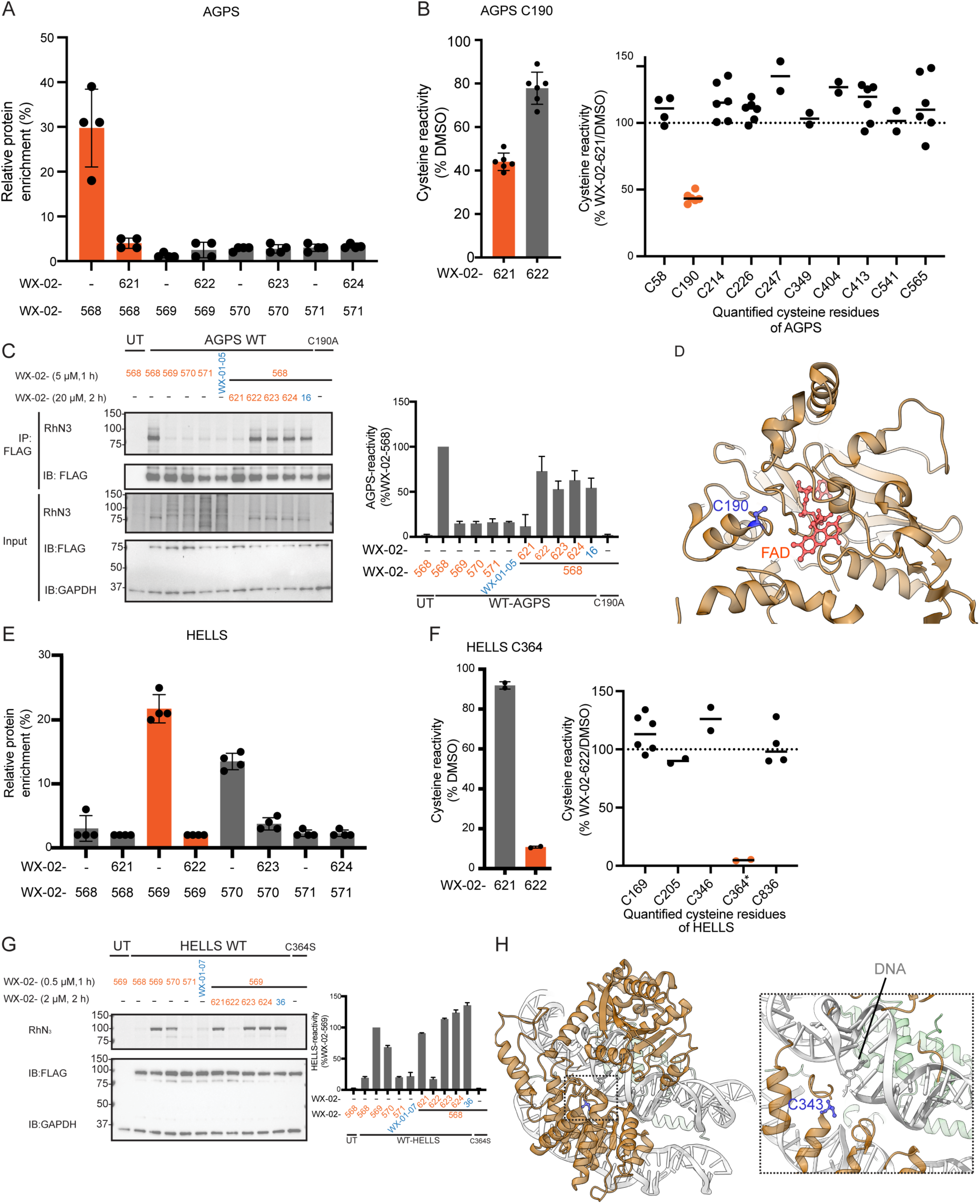
Further characterization of liganding events for representative tryptoline butynamide-preferring proteins. (A) Protein-directed ABPP data showing stereoselective enrichment of AGPS by WX-02-568 and blockade of this enrichment by WX-02-621. (B) Left, cysteine-directed ABPP data showing that AGPS is stereoselectively liganded at C190 by WX-02-621. Right, reactivity of all quantified cysteine residues in AGPS with WX-02-621. (C) Gel-ABPP data following anti-FLAG immunoprecipitation (IP) showing the stereoselective engagement of FLAG-tagged WT-AGPS, but not a C190A-AGPS mutant, by WX-02-568, and the blockade of WX-02-568-WT-AGPS interactions by pretreatment with WX-02-621 (20 μM, 2 h). (D) Crystal structure of *C. porcellus* AGPS (tan, PDB: 5ADZ, 2.00-2.20 Å). C190 (blue, corresponding to human AGPS C190) is located ∼10 Å away from the FAD co-factor (red). (E) Protein-directed ABPP data showing enantioselective enrichment of HELLS by WX-02-569 and WX-02-570 and blockade of these enrichments by WX-02-622 and WX-02-623, respectively. (F) Left, targeted cysteine-directed ABPP data showing that HELLS is stereoselecctively liganded at C364 by WX-02-622. Right, reactivity of all quantified cysteine residues in HELLS with WX-02-622. Cysteines other than C364 were quantified by untargeted cysteine-directed ABPP performed with a trypsin-digested proteome; C364 was quantified by targeted cysteine-directed ABPP performed with a LysC-digested proteome (asterisked). (G) Gel-ABPP data showing the stereoselective engagement of FLAG-tagged WT-HELLS, but not a C364S-HELLS mutant, by WX-02-569, and the blockade of WX-02-569-WT-HELLS interactions by pre-treatment with WX-02-622 (2 μM, 2 h). (H) Cryo-EM structure of an *A. thaliana* DDM1-nucleosome complex (DDM1, tan; histones, green; PDB: 7UX9, 3.20 Å)^87^ and a zoom-in look on the right showing a relative distance of 4.5 Å between the conserved DDM1 C343 (blue, corresponding to HELLS C364) and the nearest DNA strand (light gray). IP, immunoprecipitation using anti-FLAG antibodies; IB, anti-FLAG and anti-GAPDH immunoblot; UT, untransfected cells. In A, B, C, E, F, and G, data represent average values ± s.d. from two independent experiments. In C and G, gel images are representative of two independent experiments.

HELLS, or lymphoid-specific helicase (LSH), is an ATP-dependent chromatin remodeling protein involved in DNA replication, transcription, and repair.^83–85^ Protein-directed ABPP experiments revealed the enantioselective liganding of HELLS by (1*R*, 3*R*) and (1*R*, 3*S*) alkyne/parent tryptoline butynamide pairs WX-02-569/WX-02-622 and WX-02-570/WX-02-623, respectively (**Figure 3E**). The stereoprobe-liganded cysteine in HELLS was not identified in our cysteine-directed ABPP experiments, despite these experiments quantifying several cysteines in the protein (**Figure 3F**). We have previously found that cysteines liganded by electrophilic compounds may evade detection by ABPP if they reside on non-proteotypic (e.g., very large or small) tryptic peptides.^42^ Performing a second set of targeted cysteine-directed ABPP experiments with an alternative protease (Lys-C) digestion identified C364 as the site of enantiose-lective liganding in HELLS by WX-02-622 (**Figure 3F**). This cysteine is located within the primary amino acid sequence …KC364RLIRELK…, providing a clear explanation why it was detected in cysteine-directed ABPP experiments performed with Lys-C (which generated an eight amino acid peptide), but not trypsin (which generated a two amino acid peptide), as the digestion protease.

Gel-ABPP experiments confirmed that WX-02-569 and WX-02-570 each enantioselectively engaged recombinant WT-HELLS expressed in HEK293T cells, while the tryptoline acrylamide stereomatched to WX-02-569 did not react with WT-HELLS (**Figure 3G**). WX-02-569 did not engage a C364S-HELLS mutant and its interaction with WT-HELLS was stereoselectively blocked by pre-treatment of HEK293T cells with WX-02-622 in a concentration-dependent manner (IC50 = 0.3 µM), but not by the stereomatched tryptoline acrylamide WX-02-36 (IC50 > 2 µM; **Figures 3G** and **S5**). A cryo-electron microscopy (cryo-EM) structure of human HELLS has recently been reported,^86^ but the coordinates of this structure are not yet publicly available. Instead, we analyzed a cryo-EM structure of the *A. thaliana* ortholog of HELLS (DDM1; PDB: 7UX9, 3.20 Å),^87^ which indicated that the butynamide-liganded C364 is in a conserved region in close proximity to the DNA-protein interaction surface (**Figure 3H**). These data thus suggest that tryptoline butynamides like WX-02-622 could be useful chemical probes for studying the functions of HELLS as a chromatin remodeler.^88, 89^

### Tryptoline butynamides inhibit the actin maturation protease ACTMAP

Prominent among the butynamide-preferring proteins was C19orf54, or ACTMAP, which showed robust stereoselective reactivity with the (1*R*, 3*S*) alkyne/parent tryptoline butynamide pair WX-02-570/WX-02-623, but negligible reactivity with the stereomatched tryptoline acrylamides (**Figure 4A, B**). ACTMAP is a ∼45 kDa cytosolic protein identified in a haploid genetic screen to function as a protease that post-translationally cleaves the *N*-terminally acetylated methionine from β- and γ-actin, which are then reacetylated by *N*-acetyltransferase NAA80 to generate mature actins.^73^ AlphaFold predictions revealed that ACTMAP has structural similarity to bacterial cysteine proteases with C132 representing the catalytic nucleophile.^73^ While our cysteine-directed ABPP experiments did not directly identify the butynamide-reactive cysteine in ACTMAP, we confirmed the stereoselective reactivity of WX-02-570 with recombinant WT-ACTMAP, but not a C132A mutant, in HEK293T cells by gel-ABPP (**Figures 4C** and **S6A**). Other representative cysteine mutants of ACTMAP (C119A and C271A) retained reactivity with WX-02-570 (**Figure S6B**). We note that C132 is part of a long tryptic peptide (a.a. 115-154), which may explain why this site was not quantified in cysteine-directed ABPP experiments.

**Figure 4.**
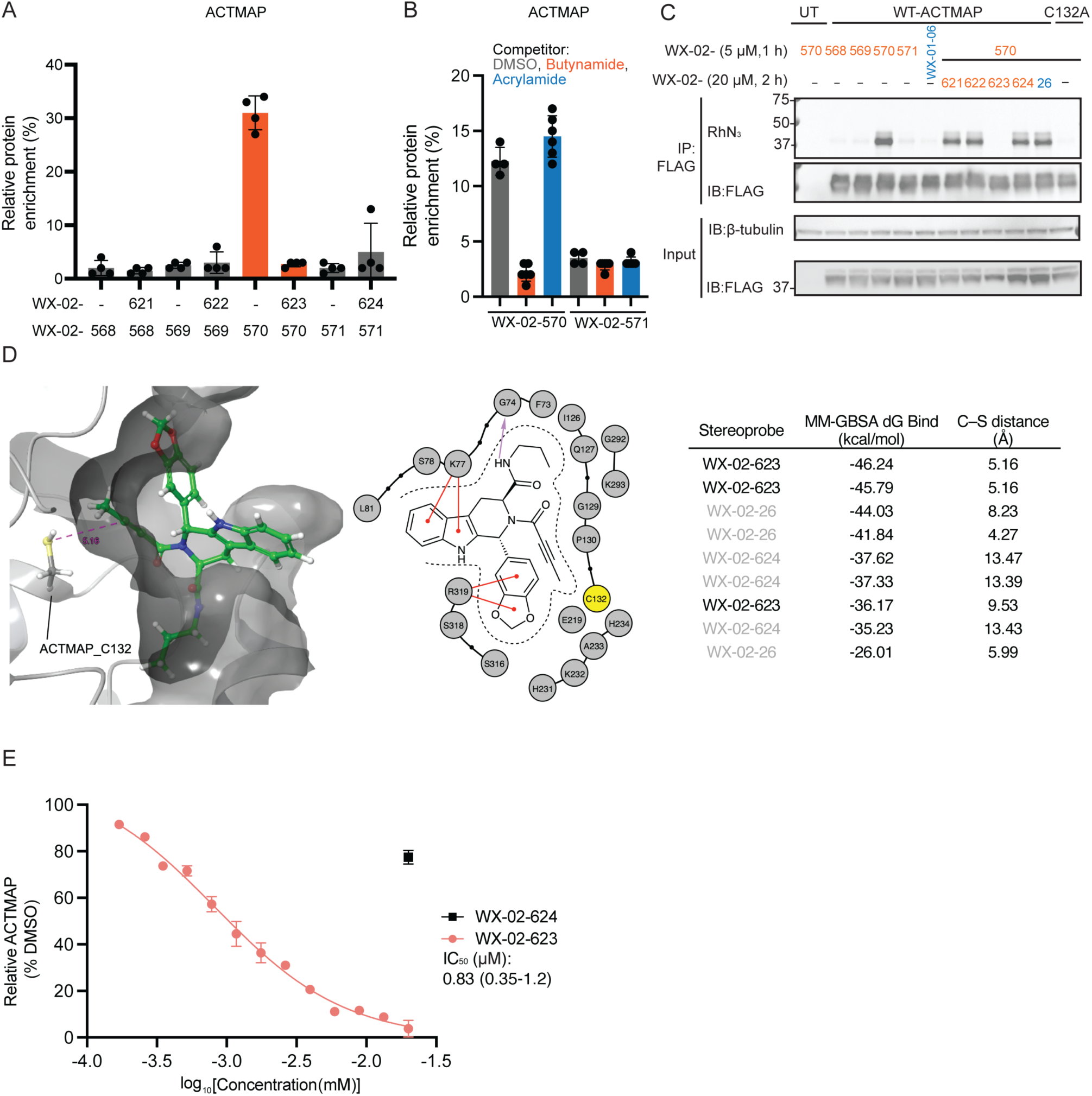
Further characterization of the butynamide-preferring protein C19orf54/ACTMAP. (A) Protein-directed ABPP data showing the stereoselective enrichment of ACTMAP by WX-02-570 and blockade of this enrichment by WX-02-623. (B) Cross-competition ABPP data showing blockade of WX-02-570 enrichment of ACTMAP by the stereomatched butynamide WX-02-623, but not the corresponding acrylamide WX-02-26. (C) Gel-ABPP data following anti-FLAG immunoprecipitation (IP) showing the stere-oselective engagement of FLAG-tagged WT-ACTMAP, but not a C132A-ACTMAP mutant, by WX-02-570 (5 µM, 1 h), and the blockade of WX-02-570-WT-ACTMAP interactions by pretreatment with WX-02-623 (20 μM, 2 h). (D) Non-covalent docking on AlphaFold2 model recapitulates the observed stereoselective liganding of ACTMAP by WX-02-623. Left, lowest-energy binding pose of WX-02-623 in the active site of ACTMAP (dG Bind -46.24 kcal/mol). The pink dashed line denotes the distance from the sulfur atom of the C132 catalytic nucleophile to the electrophilic carbon atom of WX-02-623 (5.16 Å). Image generated with Schrödinger Maestro. Middle, WX-02-623-ACTMAP interaction diagram showing ACTMAP residues located within 4 Å of any atom of the ligand. Red lines denote pi-cation interactions, purple arrows denote H-bonding interactions, dashed lines denote the protein surface. ACTMAP_C132 is highlighted in yellow. Right, table comparing docking output poses, showing MM-GBSA scores (kcal/mol) and distances from the sulfur atom of the C132 catalytic nucleophile to the electrophilic carbon atom of the ligands (C–S distance, Å). (E) Quantification of gel-ABPP data for concentration-dependent blockade of WX-02-570-WT-ACTMAP interactions by WX-02-623, from which an IC50 value of 0.83 μM (95% confidence interval of 0.35 to 1.2 μM) was calculated. See **Figures S6C** for representative gel-ABPP data. IP, immunoprecipitation using anti-FLAG antibodies; IB, anti-FLAG and anti-GAPDH im-munoblot; UT, untransfected cells. In A, B, and E, data represent average values ± s.d. from two independent experiments. In C, gel image is representative of two independent experiments.

Docking experiments performed with an AlphaFold2-generated^90–92^ structural model of ACTMAP supported the preferred interactions with WX-02-623 over the inactive enantiomer WX-02-624 and positioned the butynamide reactive group in close proximity to C132 (**Figure 4D**). We determined by gel-ABPP that pre-treatment with WX-02-623 (2 h) blocked WX-02-570 reactivity with recombinant ACTMAP in HEK293T cells with an IC50 value of 0.83 μM (95% CI of 0.35 – 1.20 µM (**Figures 4E** and **S6C**). We estimated a similar potency of engagement for WX-02-623 with endogenous ACTMAP in 22Rv1 cells by protein-directed ABPP (IC50 value ∼2 µM; **Figure S6D**). The inactive enantiomer WX-02-624 showed negligible interactions with recombinant or endogenous ACTMAP at 20 µM test concentration (**Figures 4E** and **S6C, D**).

We next tested whether WX-02-623 inhibited ACTMAP activity in cancer cells. We first confirmed, as has been shown previously,^73^ that genetic disruption of ACTMAP by CRISPR/Cas9 gene editing resulted in substantial accumulation of immature actin as measured by Western blotting with an antibody recognizing the *N*-terminus of β-actin^73^ (**Figure S7A**). A similar effect was observed in 22Rv1 cells treated with WX-02-623 (1-20 µM, 48 h), which produced a concentration-dependent increase in immature β-actin (**Figure 5A**). In contrast, we did not detect accumulation of immature β-actin in WX-02-624-treated cells (**Figure 5A**). A time-course study revealed that immature β-actin could be detected in cells as early as 4 h post-treatment with WX-02-623 (**Figure 5B**).

**Figure 5.**
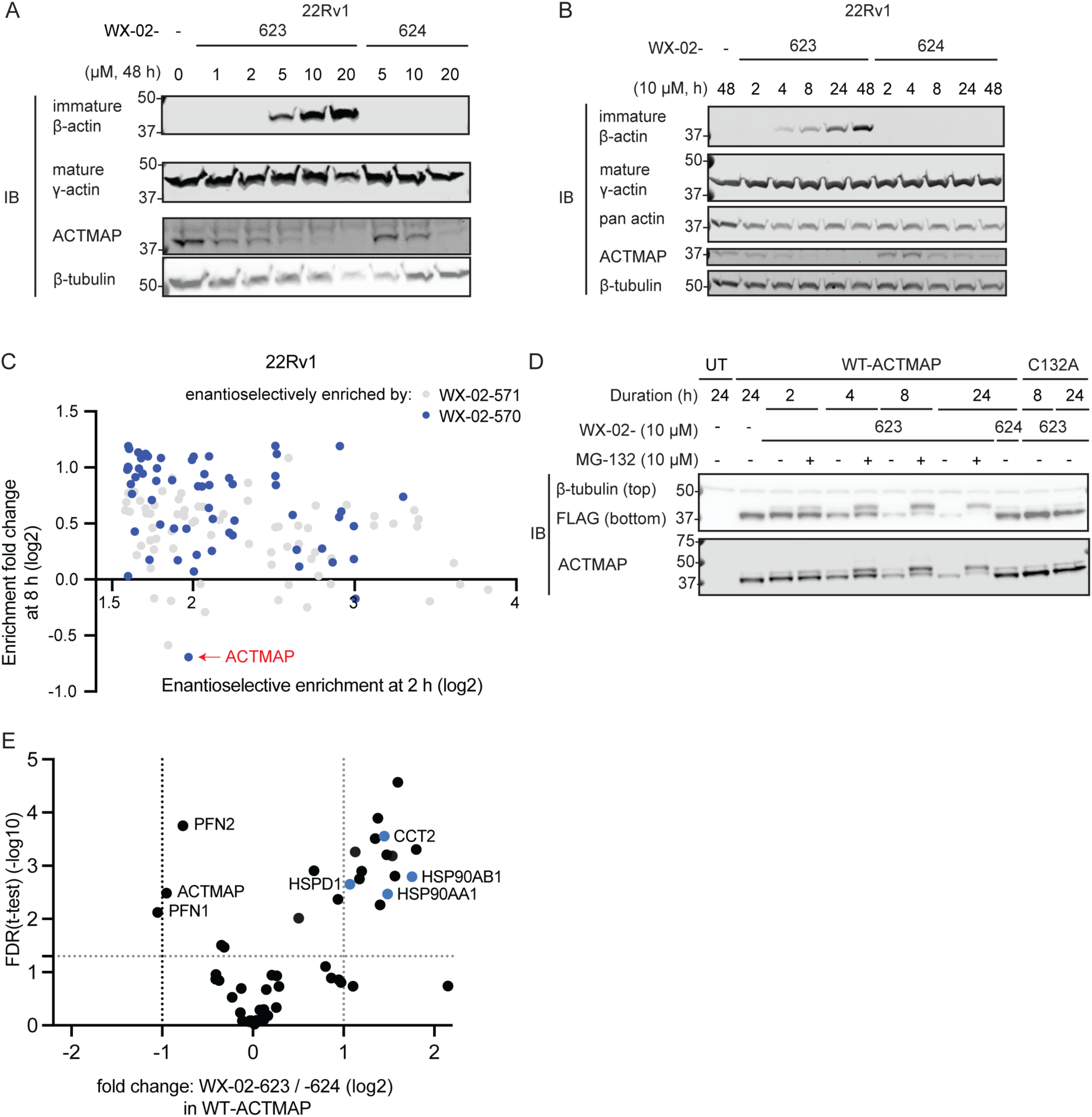
Characterization of the effects of tryptoline butynamide WX-02-623 on ACTMAP function in cells. (A, B) Western blots showing concentration- (A) and time- (B) dependent effects of WX-02-623 or WX-02-624 on immature β-actin accumulation in 22Rv1 cells. Note that mature γ–actin was measured instead of mature β-actin because antibodies specific for mature β-actin are lacking, as detailed previ-ously.^73^ (C) Scatter plot showing the enantioselective enrichment values of proteins at 2 h (x-axis) versus the fold-change in enrichment for these proteins with the preferred stereoprobe at 2 vs 8 h treatment (y-axis) for 22Rv1 cells treated with WX-02-570 or WX-02-571. (D) Western blot showing WT-ACTMAP, but not a C132A mutant, is enantioselective degraded by WX-02-623 in a time- and proteasome (MG132)-dependent manner. HEK293T cells recombinantly expressing FLAG-tagged WT-ACTMAP or C132A-ACTMAP were treated with WX-02-623 or WX-02-624 (10 μM) for the indicated times with or without MG-132 (10 μM). (E) Volcano plot of anti-FLAG immunoprecipitation-mass spectrometry (IP-MS) data comparing protein abundance measurements from WT-ACTMAP-expressing HEK293T cells treated with WX-02-623 or WX-02-624 (10 μM, 3 h). Chaperone/Co-chaperone proteins are highlighted in blue. IB, anti-FLAG, anti-immature-β-actin, anti-γ-actin, anti-pan-actin, anti-β-tubulin, and anti-ACTMAP im-munoblot; UT, untransfected cells. In A, B, and D, blot images are representative of at least two independent experiments. In C, data are average values from two independent experiments. In E, data represent average values from three independent experiments.

In cells treated with WX-02-623, we also observed an apparent decrease in the ACTMAP protein itself by Western blotting that was stereoselective at test concentrations below 20 µM (**Figure 5A, B**). This result suggested that liganding by WX-02-623 might lead to the degradation of ACTMAP. Consistent with this hypothesis, we performed a time-dependent protein-directed ABPP experiment.^34^ which revealed that among the proteins that were enantioselectively enriched by WX-02-570, ACTMAP uniquely showed a loss of enrichment at 8 vs 2 h (**Figures 5C** and **S7B**). We observed a similar time-dependent and stereoselective loss of recombinant WT-ACTMAP, but not the C132A-ACTMAP mutant, in HEK293T cells treated with WX-02-623 (**Figure 5D**). The WX-02-623-dependent decrease in WT-ACTMAP was blocked by the proteasome inhibitor MG132, which also led to accumulation of an upward band-shifted form of WT-ACTMAP that may represent a covalent adduct with WX-02-623 (**Figure 5D**). The Neddylation inhibitor MLN4924 (1 µM) did not block WX-02-623-dependent loss of WT-ACTMAP (**Figure S7C**), suggesting that a cullin-RING-E3 ligase is not involved in the degradation process. Finally, we found in co-immunoprecipitation (IP)-MS experiments that WX-02-623, but not WX-02-624, promoted the binding of ACTMAP to several chaperone proteins – HSP90AA1, HSP90AB1, and CCT2 (**Figure 5E** and **Dataset S1**) – possibly indicating that reactivity with WX-02-623 destabilized the folded stated of ACTMAP to promote binding to chaperones.^93^ These IP-MS experiments also revealed stereoselective reductions in the enrichment of established ACTMAP-interacting proteins, such as profilin 1 and 2 (PFN1/2)^73, 94, 95^ (**Figure 5E**), in WX-02-623-treated cells, but the concurrent decrease in ACTMAP itself (**Figure 5E**) suggests that these effects may be indirectly related to ligand-induced degradation of ACTMAP.

## Conclusion

The comparative ligandability maps established herein have revealed that the tryptoline butynamides, despite showing lower intrinsic and proteomic reactivity than tryptoline acrylamides, preferentially liganded several proteins in human cells. Such liganding events, which we confirmed for representative proteins by recombinant expression, may reflect instances where the butynamide is better oriented in a pocket to react with a proximal cysteine or where the microenvironment of the pocket preferentially activates the butynamide.^75^ Regardless of the precise mechanistic basis for butynamide-preferring liganding events, they were found on diverse proteins from distinct structural and functional classes in our ABPP experiments, supporting the broad potential of the butynamide as a source of covalent chemical probes. Considering further that few butynamides have been examined to date for their proteome-wide reactivity,^17, 41^ we anticipate that ABPP investigations of more structurally diverse sets of compounds should identify additional proteins that react with this electrophile. We also note that, even for proteins showing similar degrees of liganding with acrylamides and butynamides, the latter electrophile may be preferred for probe optimization due to its lower intrinsic reactivity and affordance of an additional site for chemical derivatization (the butynamide methyl group).

The identification of tryptoline butynamides that stereoselectively inhibit the cysteine protease ACTMAP should offer valuable chemical tools to complement genetic methods for studying the post-translational modification of actin in various biological settings, akin to the utility of covalent inhibitors and activity-based probes targeting other types of cysteine proteases.^96–99^ ACTMAP-knockout mice are viable, but exhibit decreased muscle strength due to the accumulation of immature actin and shortening of sarcomeric actin microfilaments.^73^ The actin cytoskeleton also plays important roles in cell proliferation and migration, where there is interest in identifying ways to selectively perturb actin microfilament dynamics in cancer cells.^100–102^ Exploring the impact of ACTMAP inhibitors in different cancer settings could accordingly represent an interesting area for future investigation. We do not yet understand how covalent liganding by tryptoline butynamides leads to degradation of ACTMAP in cells, but our data suggest a proteasome-dependent process that may involve E3 ligases outside of the cullin-Ring E3 ligase family. Genetic screens^103, 104^ may help to identify the proteins that mediate tryptoline butynamide-induced degradation of ACTMAP. More generally, it is important to emphasize that additional studies are needed to determine if and how the other butynamide-preferring liganding events identified herein affect the functions of proteins such as AGPS and HELLS. The stereoselectivity and site-specificity of these tryptoline butynamide-protein interactions should provide useful chemical (inactive enantiomer) and genetic (stereoprobe-resistant cysteine mutant) controls for such biological studies.^23, 33, 34, 41–43^

## ASSOCIATED CONTENT

### Supporting Information

The Supporting Information is available free of charge on the ACS Publications website.

Supplementary Figures (PDF)

Supporting Data Set 1. A Global Ligandability Map of Tryptoline Butynamide Stereoprobes Identifies Covalent Inhibitors of the Actin Maturation Protease ACTMAP (xlsx)

Supporting Table 1. GSH reactivity assay (doc)

Supporting Document (PDF): Reagents, methods, and synthetic chemistry

## AUTHOR INFORMATION

### Authors

Yijun Xiong - Department of Chemistry, Scripps Research, La Jolla, California 92037, United States

Christopher J. Reinhardt - Department of Chemistry, Scripps Research, La Jolla, California 92037, United States

Tracey Nguyen - Vividion Therapeutics, San Diego, California 92121, United States

Melissa A. Hoffman - Vividion Therapeutics, San Diego, California 92121, United States

Gabriel M. Simon - Vividion Therapeutics, San Diego, California 92121, United States

## Notes

The authors declare the following competing financial interest(s): Drs. Hoffman, Nguyen, and Simon are employees to Vividion Therapeutics. Dr. Cravatt is a founder and scientific advisor to Vividion Therapeutics. Vividion is a biotechnology company interested in developing small-molecule medicines.

## ACKNOWLEDGMENTS

The authors thank Prof. Howard C. Hang, Prof. Michael A. Erb, Prof. Michalina Janiszewska, Prof. Ryan A. Shenvi, Dr. Zachary E. Potter, Dr. Timothy Ware, Dr. Wieland Goetzke, Dr. Sang Joon Won, Dr. Haoxin Li, Garrett L. Lindsey, Rachel E. Hayward, Xuanmeng Luo, and Elva Ye for their general assistance and insights. The authors thank Dr. Melissa Dix, Dr. Daisuke Ogasawara, and Sabrina Barbas for assistance with proteomics experiments. We also thank Dr. Thijn R. Brummelkamp (Netherlands Cancer Institute) for providing antibodies that detect immature actins. We thank Dr. Xuedong Liu and Dr. Bing Chen (WuXi AppTec) for small-molecule synthesis and Quynh Nguyen Wong, Jillian Smith, Jason Lee, Catherine Chiang, and Dr. Brandon Orzolek (Scripps Automated Synthesis Facility) for support with high-resolution mass spectrometry. Unless otherwise noted, protein structure figures were generated with UCSF ChimeraX, developed by the Resource for Biocomputing, Visualization, and Informatics at the University of California, San Francisco, with support from National Institutes of Health R01-GM129325 and the Office of Cyber Infrastructure and Computational Biology, National Institute of Allergy and Infectious Diseases.

---

Insert Table of Contents artwork here

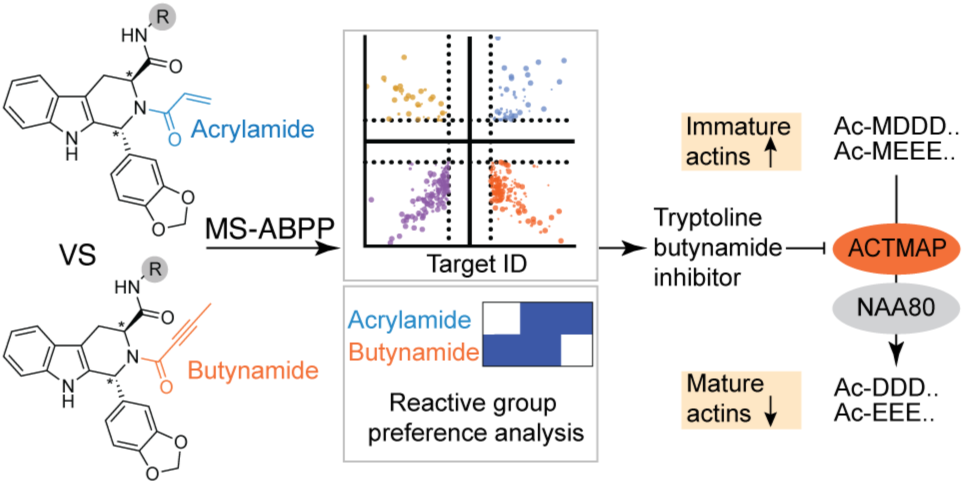

**Figure S1.**
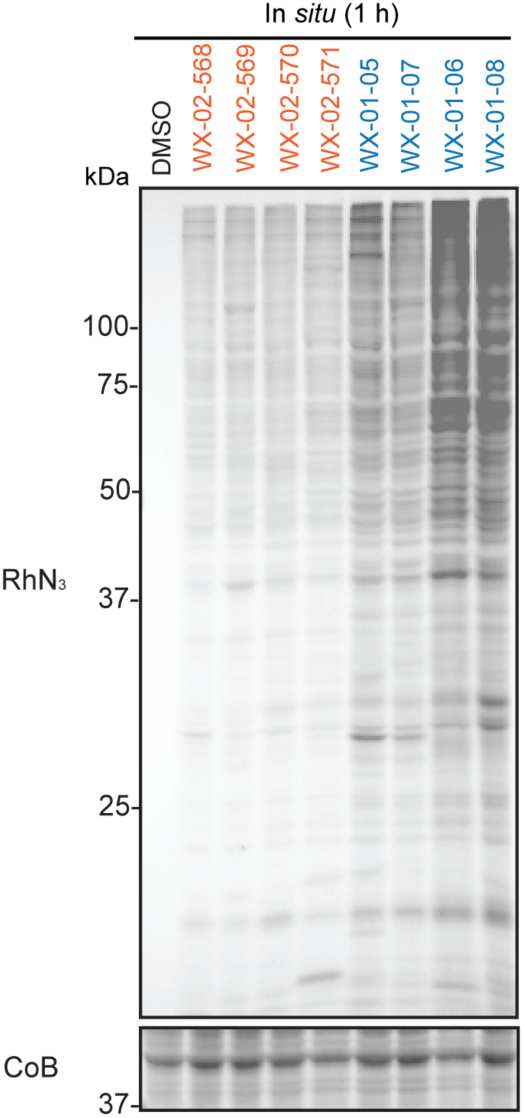
Gel-ABPP data for Ramos cells treated with alkyne stereoprobes (10 µM) for 1 h, followed by lysis and visualization of stereoprobe-reactive proteins by copper-catalyzed azide-alkyne cycloaddition (CuAAC) conjugation^76, 77^ to an azide-rhodamine (RhN3) reporter group, SDS-PAGE, and in gel fluorescence scanning.^105^ CoB, Coomassie blue staining. The names of tryptoline butynamides and tryptoline acrylamides are shown in orange and blue font, respectively. Data correspond to the gel-ABPP experiment shown in Figure 1D (exposure time 240 s), here with shorter exposure time (120 s).

**Figure S2.**
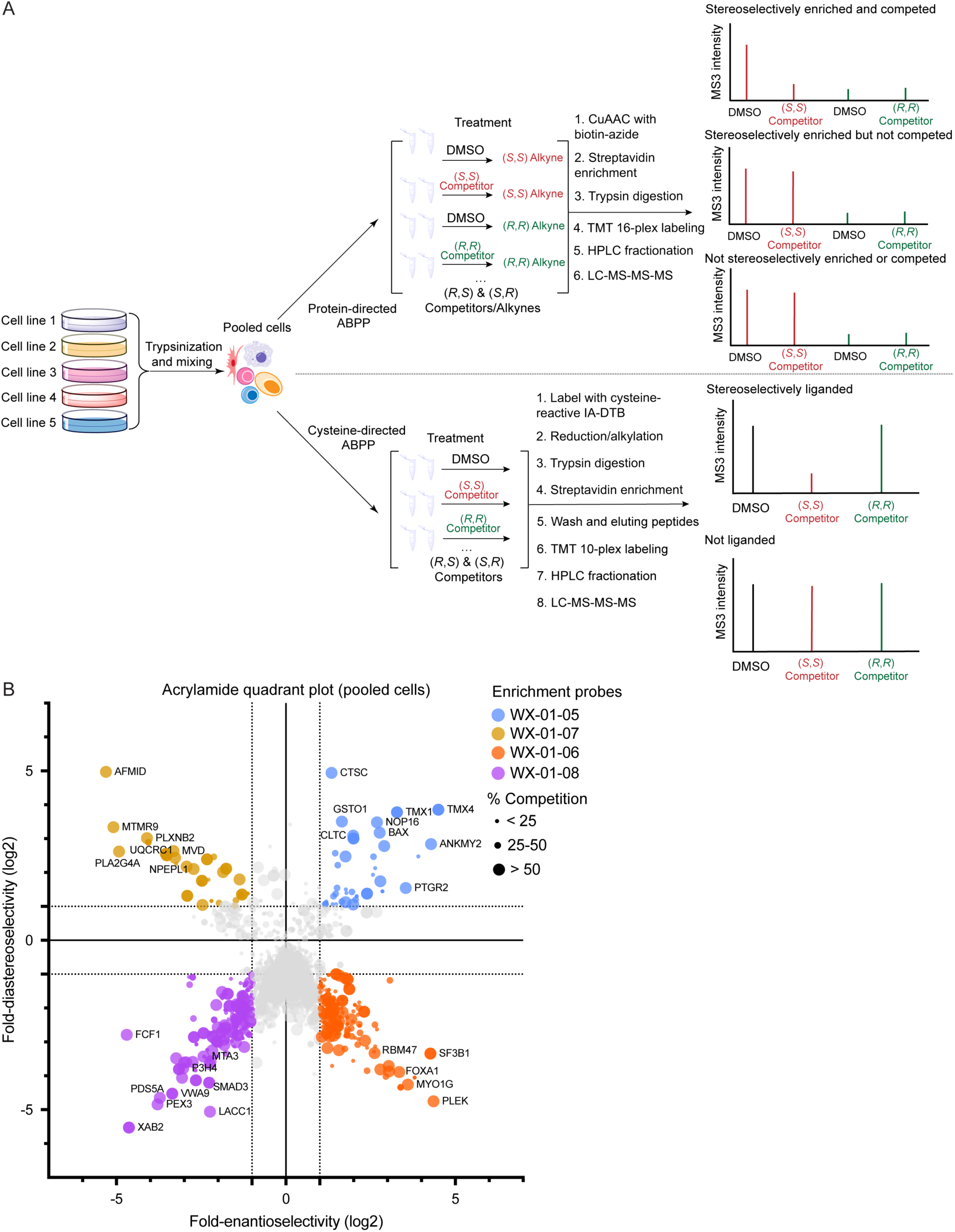
Mapping proteins preferentially liganded by tryptoline butynamides and acrylamides by protein-directed ABPP. (A) General workflow for protein-directed and cysteine-directed ABPP: Ramos, 22Rv1, UACC-257, U-2 OS, and Hep G2 were cultured separately and pooled on the day of treatment based on equal protein content predetermined by bicinchoninic acid assay (BCA) assay. All cell treatments were performed in a pooled suspension state. Protein-directed ABPP: pretreatment with DMSO or parent competitor stereoprobes (20 µM) for 2 h, followed by stereomatched alkyne stereoprobe (5 µM) for 1 h; stereoselective enrichment of proteins by alkyne stereoprobes and blockade of enrichment by stereomatched non-alkyne competitor stereoprobes were determined by multiplexed (tandem mass tagging, TMT^16plex^) MS-based proteomics.^41, 42^ Cysteine-directed ABPP: treatment with parent stereoprobes (20 µM) for 3 h; stereoprobe reactivity with cysteines was determined by multiplexed (TMT^10plex^) MS-based proteomics.^70^ Mock data on the right show examples of liganded proteins (protein-directed ABPP) and cysteines (cysteine-directed ABPP). (B) Quadrant plots highlighting stereoselectively liganded proteins for each stereoconfiguration of tryptoline acrylamides as determined by protein-directed ABPP. Enantioselectivity (x axis) is the ratio of enrichment for one stereoisomer versus its enantiomer, and diastereoselectivity (y axis) is the ratio of enrichment of one stereoisomer versus the average of its two diastereomers.

**Figure S3.**
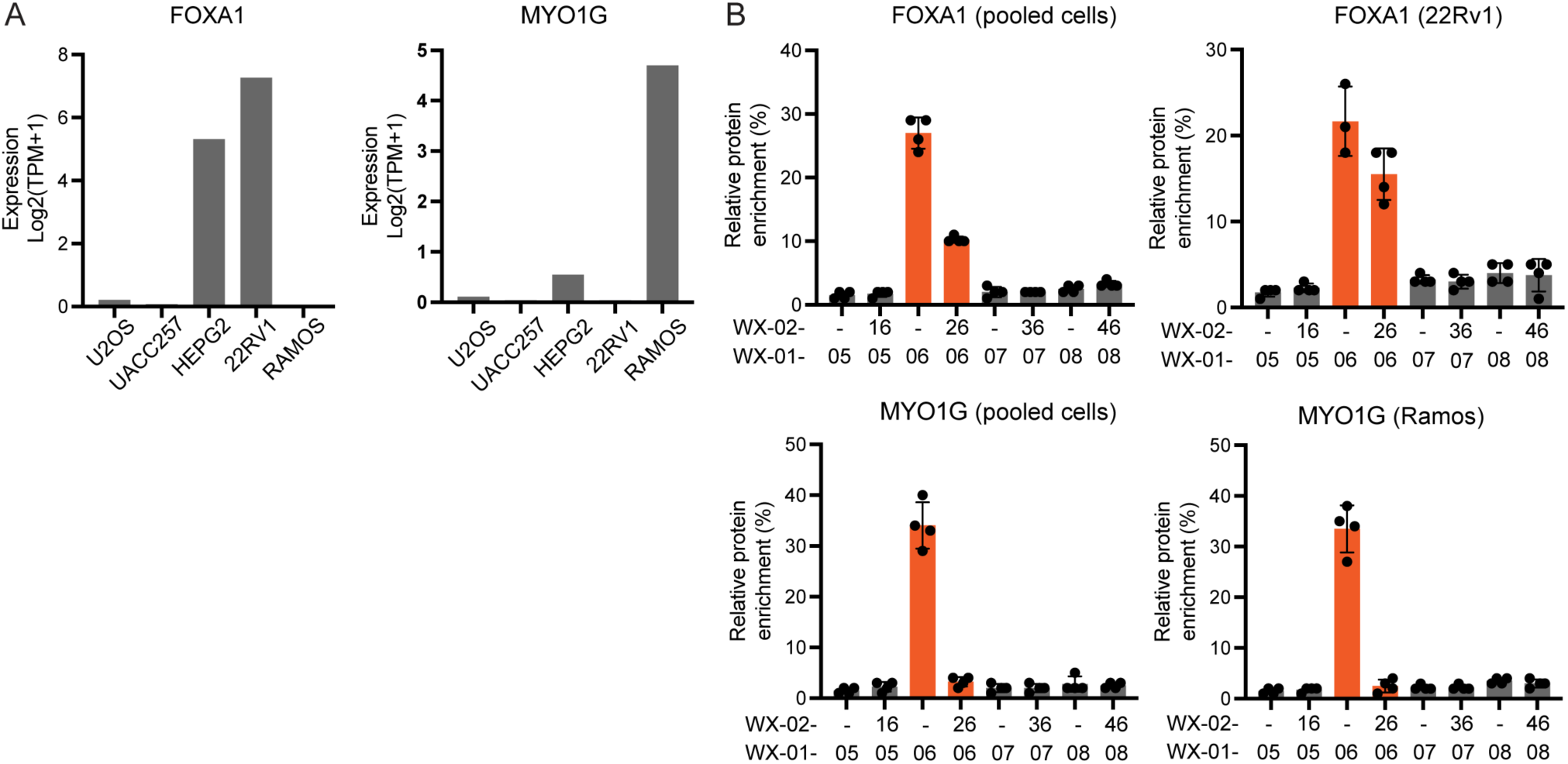
Quantification of proteins with cell line-restricted expression in protein-directed ABPP experiments performed with pooled cell lines. (A) Gene expression levels for two cell line-restricted proteins FOXA1 (left) and MYO1G (right) (RNA-seq data extracted from DepMap, 2023Q3 release).^106^ (B) Protein-directed ABPP data for pooled cells vs individual cell lines^42^ showing similar stereoprobe liganding profiles for FOXA1 and MYO1G. For B, data represent average values from four independent experiments; error bars represent s.d. values.

**Figure S4.**
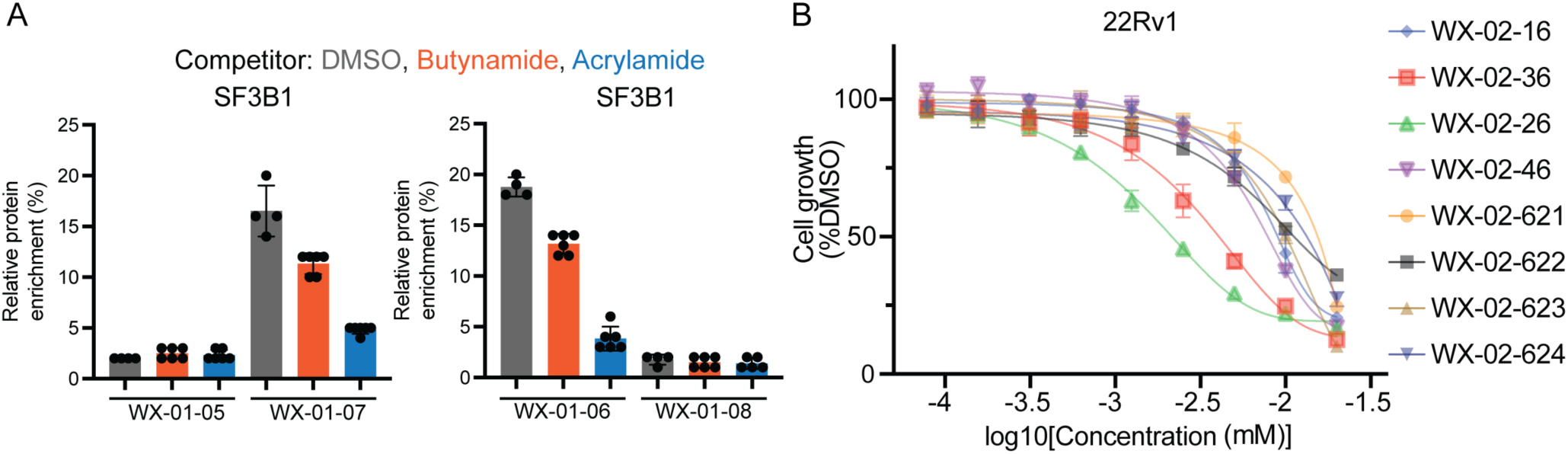
Tryptoline butynamides show attenuated engagement of the spliceosome factor SF3B1 and negligible stereoselective anti-proliferative effects. (A) Cross-competition protein-directed ABPP data for SF3B1, where enrichment was performed with alkyne tryptoline acrylamides. Pooled cell lines were treated with DMSO or parent tryptoline butynamides or acrylamides (20 µM) for 2 h followed by the stereomatched alkyne tryptoline acrylamide (5 µM) for 1 h. Workflow as shown in Figure 2B. (B) Con-centration-dependent effects of tryptoline acrylamides and butynamides on the proliferation of 22Rv1 cells. Cells were treated with the indicated concentrations of compounds or DMSO, and, after 3 d, cell proliferation was determined by CellTiter-Glo (CTG). For A, data represent average values from four independent experiments. For B, data represent average values from three technical replicates from one experiment and are normalized to DMSO-treated cells. For A and B, error bars represent s.d. values.

**Figure S5.**
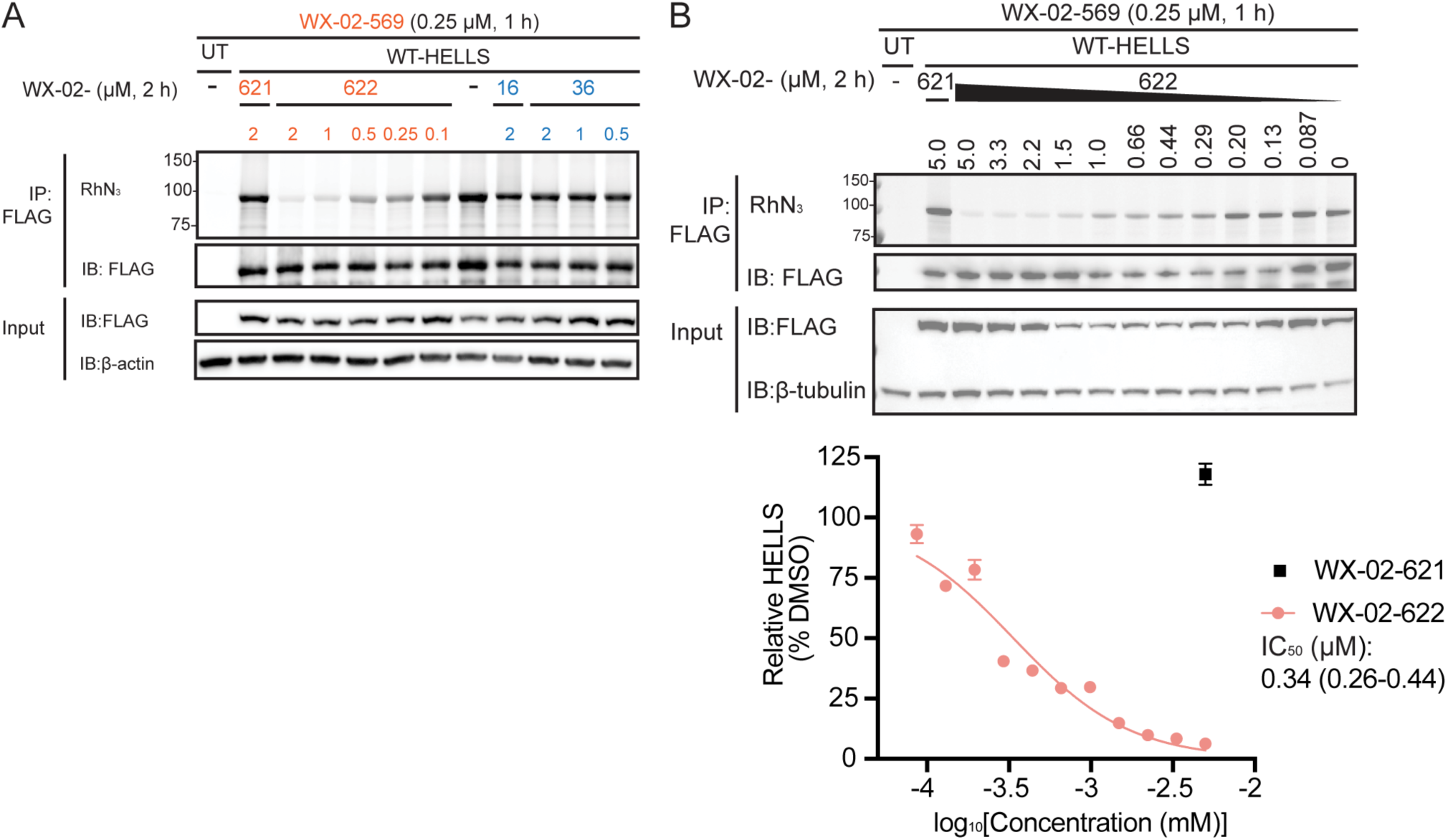
Further characterization of the butynamide-preferring protein HELLS. (A) Gel-ABPP data following anti-FLAG immunoprecipitation (IP) showing the stereoselective engagement of FLAG-tagged WT-HELLS by WX-02-569 (0.25 µM, 1 h), and the concentration-dependent blockade of WX-02-569-WT-HELLS interactions by pretreatment with tryptoline butynamide WX-02-622, but not with tryptoline acrylamide WX-02-36 (2 h). (B) Top, Gel-ABPP data following anti-FLAG IP showing the stereoselective engagement of FLAG-tagged WT-HELLS by WX-02-569 (0.25 µM, 1 h), and the concentration-dependent blockade of WX-02-569-WT-HELLS interactions by pretreatment with tryptoline butynamide WX-02-622 (2 h, 12-point concentration curve). Gel image is representative of two independent experiments. Bottom, quantification of concentration-dependent blockade of WX-02-569-WT-HELLS interactions by WX-02-622, from which an IC50 value of 0.34 μM (95% confidence interval of 0.26 to 0.44 μM) was calculated. Data are shown as average values ± s.d. from two independent experiments. IB: anti-FLAG, anti-β-actin, and anti-β-tubulin immunoblot; UT, untransfected cells.

**Figure S6.**
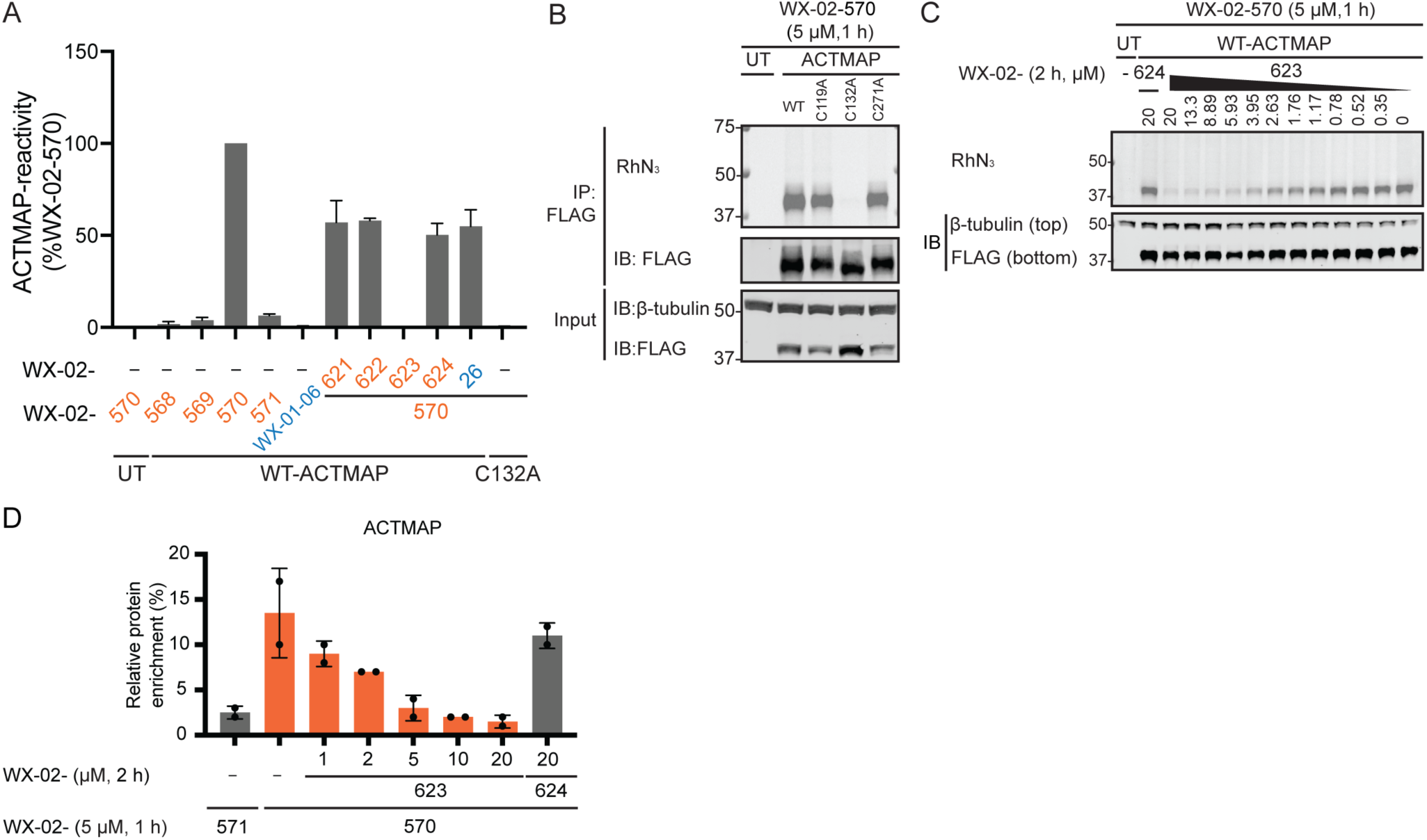
Further characterization of the butynamide-preferring protein C19orf54/ACTMAP. (A) Quantification of gel-ABPP data following anti-FLAG immunoprecipitation (IP) showing the stereoselective engagement of FLAG-tagged WT-ACTMAP, but not a C132A-ACTMAP mutant, by WX-02-570 (5 µM, 1 h), and the blockade of WX-02-570-WT-ACTMAP interactions by pretreatment with WX-02-623 (20 μM, 2 h) (see representative gel-ABPP data in Figure 4C). (B) Gel-ABPP showing the engagement of WT-ACTMAP and C119A- and C132A-ACTMAP mutants, but not a C132A-ACTMAP mutant, by WX-02-570 (5 μM, 1 h). (C) Gel-ABPP data showing concentration-dependent blockade of WX-02-570-WT-ACTMAP interactions by WX-02-623; image is representative of two independent experiments used to calculate and IC50 value for WX-02-623 (see Figure 4E). (D) Concentration-dependent engagement of endogenous ACTMAP by WX-02-623 measured by protein-directed ABPP. 22Rv1 cells were treated with the indicated concentrations of WX-02-623 or WX-02-624 for 2 h followed by WX-02-570 and WX-02-571, respectively (5 μM, 1 h) and analysis by protein-directed ABPP. IP, anti-FLAG immunoprecipitation; IB, anti-FLAG and anti-β-tubulin immunoblot; UT, untransfected cells. In A, data are average values ± s.d. from two independent experiments. In B and C, data are representative of two independent experiments. In D, Data are average values ± s.d. from two independent experiments.

**Figure S7.**
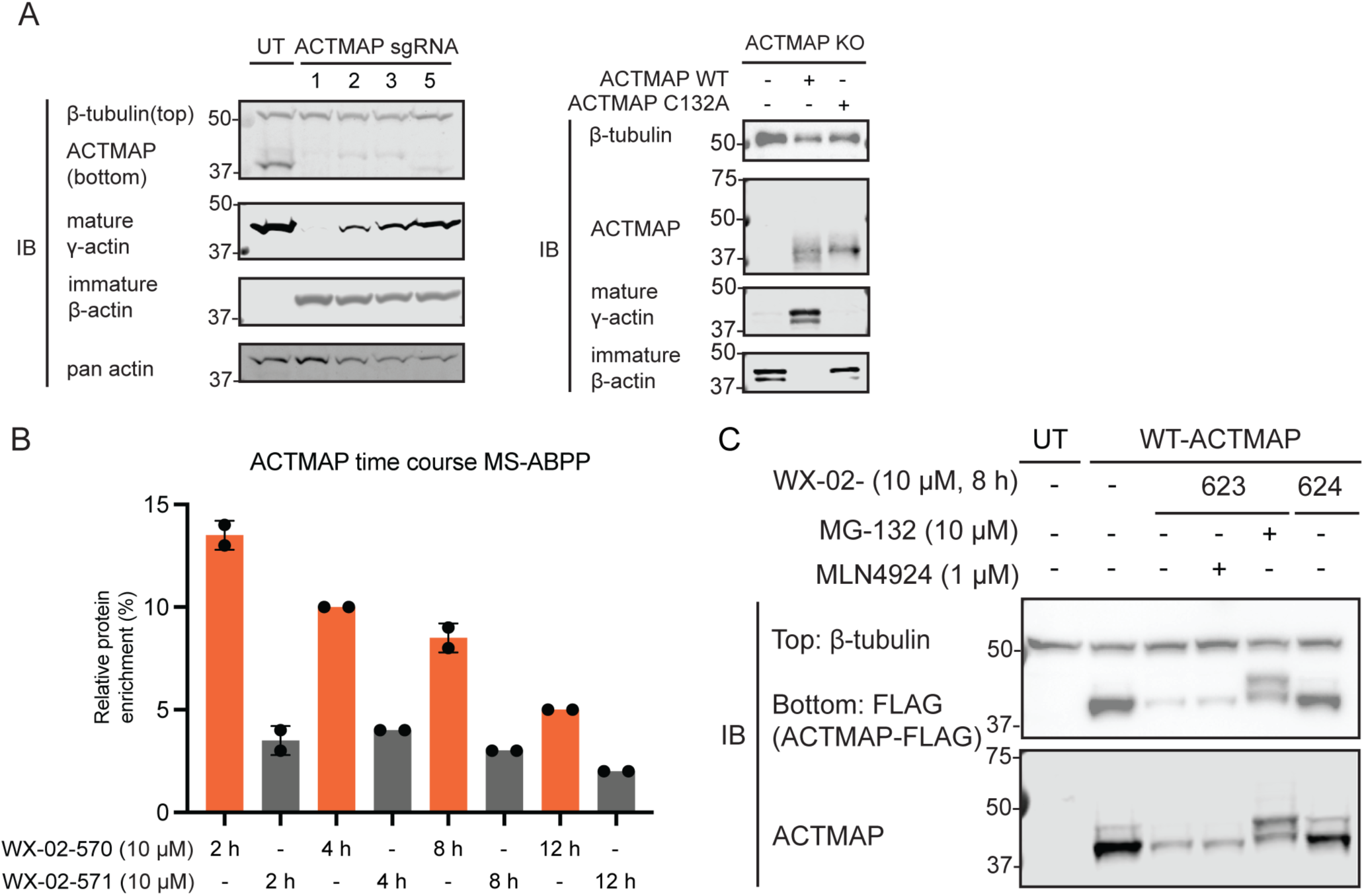
Characterization of the effects of tryptoline butynamide WX-02-623 on ACTMAP function in cancer cells. (A) Left, Western blots showing that the CRISPR/Cas9-mediated knockout (KO) of ACTMAP leads to accumulation of immature actin. Right, Western blots showing that, under ACTMAP-KO conditions, reconstitution of WT- ACTMAP, but not a C132A mutant, leads to depletion of immature actin. Due to lack of mature β-actin specific antibody, an antibody for mature γ-actin was used to measure actin maturation as previously reported.^73^ (B) Protein-directed ABPP data showing evidence for the time-dependent degradation of ACTMAP. 22Rv1 cells were treated with alkyne tryptoline butynamide WX-02-570 or WX-02-571 (10 µM) for the indicated times. (C) Western blots showing ACTMAP degradation is rescued by co-treatment with MG-132, but not MLN4924. HEK293T cells recombinantly expressing FLAG epitope-tagged ACTMAP were co-treated with WX-02-623 (10 µM) and MLN4924 (1 µM) or MG-132 (10 µM) for 8 h. IB, anti-FLAG, anti-immature-β-actin, anti-γ-actin, anti-β-tubulin, and anti-ACTMAP immunoblot; UT, untransfected cells. In A and C, gel images are representative of two independent experiments. In B, data are average values ± s.d. from two independent experiments.

**Table S1.** Glutathione reactivity values for tryptoline acrylamide and butynamide stereoprobes. Rates (M^-1^•s^-1^) of glutathione consumption by stereoprobes at 2 and 6 h. ND, not determined. Acrylamide data are from a previous publication.^42^

| Tryptoline acrylamides | $M^{-1}s^{-1}$ | Time (h) | Average $M^{-1}s^{-1}$ | Tryptoline butynamides | $M^{-1}s^{-1}$ | Time (h) | Average $M^{-1}s^{-1}$ | Stereocon-figuration |
| --- | --- | --- | --- | --- | --- | --- | --- | --- |
| WX-02-16 | 0.09 | 6 | 0.09 | WX-02-621 | 0.023 | 6 | 0.023 | 1*S*, 3*S* |
|  | 0.09 | 2 |  |  | ND | 2 |  |  |
| WX-02-26 | 0.12 | 6 | 0.13 | WX-02-623 | 0.043 | 6 | 0.057 | 1*R*, 3*S* |
|  | 0.13 | 2 |  |  | 0.071 | 2 |  |  |
| WX-02-36 | 0.08 | 6 | 0.08 | WX-02-622 | 0.026 | 6 | 0.028 | 1*R*, 3*R* |
|  | 0.08 | 2 |  |  | 0.029 | 2 |  |  |
| WX-02-46 | 0.10 | 6 | 0.10 | WX-02-624 | 0.05 | 6 | 0.054 | 1*S*, 3*R* |
|  | 0.10 | 2 |  |  | 0.058 | 2 |  |  |

