## Supplementary Methods for "A Global Ligandability Map of Tryptoline Butynamide Stereoprobes Identifies Covalent Inhibitors of the Actin Maturation Protease ACTMAP"

**Table of Contents**


| | |
|---|---|
| **Reagents and materials** | **S3** |
| **Methods** | **S5** |
| **Synthesis and Characterization of Novel Compounds** | **S12** |
| **References** | **S28** |

### Reagents and materials

**Table S2:** Reagents and materials used in this study.

| Reagents/materials | Source | Catalog number |
|---|---|---|
| DMEM (Dulbecco's Modified Eagle's Medium) | Corning | 15-013-CV |
| RPMI 1640 1× | Corning | 15-040-CV |
| Desthiobiotin polyethyleneoxide iodoacetamide (IA-DTB) | Santa Cruz Biotechnology | sc-300424 |
| Biotin-PEG4-azide | BroadPharm | BP-22119 |
| Urea | Fisher Scientific | M1084871000 |
| Iodoacetamide | Sigma-Aldrich | I1149 |
| Dithiothreitol (DTT) | Fisher Bioreagents | BP172 |
| Tris(2-carboxyethyl)phosphine HCl (TCEP) | Biosynth | FT01756 |
| Sequencing Grade Modified Trypsin | Promega | V5111 |
| Streptavidin Agarose Resin | Fisher Scientific | 20353 |
| High Capacity Streptavidin Agarose | Fisher Scientific | 20361 |
| Micro Bio-Spin column | Bio-Rad | 7326204 |
| NonidetTM P40 substitute (IgepalTM CA-630) | USB Corporation | 19628 |
| Triethylammonium bicarbonate buffer pH 8.5 | Sigma-Aldrich | T7408 |
| TMT10plex™ Isobaric Label Reagent Set | ThermoFisher Scientific | 90406 |
| TMTpro™ 16plex Label Reagent Set | ThermoFisher Scientific | A44520 |
| Ammonium bicarbonate | Sigma-Aldrich | 09830 |
| Hydroxylamine solution 50 wt. % in water | Sigma-Aldrich | 467804 |
| Formic acid, ~98%, for mass spectrometry | Honeywell Fluka | 94318-250ML-F |
| Methanol LC-MS Grade | Fisher Scientific | A452SK-4 |
| Acetonitrile Optima™LC/MS Grade | Fisher Scientific | A955 |
| Water Optima™ LC/MS Grade | Fisher Scientific | W64 |
| Protein LoBind Tubes 1.5 mL | Fisher Scientific | 13-698-794 |
| Protein LoBind Tubes 2.0 mL | Fisher Scientific | 13-698-795 |
| Clear 384 Well Polystyrene Plates With F-Bottom | Greiner Bio | 781101 |
| DMSO | Corning | 25-950-CQC |
| Puromycin | Invivogen | ant-pr-1 |
| Blasticidin | Invivogen | ant-bl-1 |
| EPPS | Millipore | E0276 |
| Sodium Dodecyl Sulfate (SDS) Biotechnology grade, 1kg | BioPioneer | C0065 |
| Glycerol | Fisher Scientific | BP229-4 |
| Chloroform | Fisher Scientific | C297-4 |
| Tris(benzyltriazolylmethyl)amine (TBTA) | TCI | T2993 |
| Fetal Bovine Serum, Dialyzed | Omega Scientific | FB-03 |

| | | |
|---|---|---|
| One Shot Stbl3 Chemically Competent E. coli | Life Technologies | C737303 |
| cOmplete ULTRA Tablets, Protease Inhibitor Cocktail | Roche | 5892970001 |
| NuPAGE LDS Sample Buffer (4×) | Invivogen | NP0008 |
| 2-Mercaptoethanol, 99% | Acros Organics | 125472500 |
| SuperSignal™ West Pico PLUS Chemiluminescent Substrate | ThermoFisher Scientific | 34580 |
| Q5® Site-Directed Mutagenesis Kit | New England Biolabs | E0552S |
| Pierce™ Anti-DYKDDDDK Magnetic Agarose | ThermoFisher Scientific | A36797 |
| FuGENE® HD Transfection Reagent | Promega | E2311 |
| Lenti-X™ Concentrator | Takara Bio | 631231 |
| Triton™ X-100 | Sigma-Aldrich | T8787 |
| PEI MAX® - Linear Polyethylenimine Hydrochloride (MW 40,000) | Polysciences | 24765 |
| Gateway™ LR Clonase™ II Enzyme mix | Invitrogen | 11791020 |
| BsmBI-v2 | New England Biolabs | R0739S |
| InstantBlue® Coomassie Protein Stain (ISB1L) | abcam | ab119211 |
| Power Blotter Select Transfer Stack, nitrocellulose, mini | ThermoFisher Scientific | PB3210 |
| Power Blotter Select Transfer Stack, nitrocellulose, regular | ThermoFisher Scientific | PB3310 |
| Power Blotter Select Transfer Stack, PVDF, mini | ThermoFisher Scientific | PB5210 |
| Power Blotter Select Transfer Stack, PVDF, regular | ThermoFisher Scientific | PB5310 |
| GlutaMAX™ Supplement | Fisher Scientific | 35050061 |
| Pierce™ BCA Protein Assay Kits | ThermoFisher Scientific | 23225 |
| Trypsin-EDTA (0.25%), phenol red | Gibco™ | 25200114 |
| Rabbit Anti-immature beta actin antibody | Gift from Dr. Thijn Brummelkamp | PMID: 36173861 |
| Mouse Monoclonal Anti-GAPDH HRP-conjugated antibody | Proteintech | HRP-60004 |
| Mouse Monoclonal Anti-FLAG(R) M2-HRP antibody | Sigma-Aldrich | A8592 |
| Mouse Monoclonal Anti-gamma Actin antibody (2A3) | abcam | ab123034 |
| Rabbit Polyclonal Anti-C19orf54/ACTMAP antibody | Sigma-Aldrich | HPA059043 |
| Rabbit Monoclonal Anti-pan-actin antibody (D18C11) | Cell Signaling | #8456 |
| Rabbit Monoclonal Anti-beta tubulin (2O13) recombinant HRP-conjugated antibody | Proteintech | HRP-80713 |
| Mouse Monoclonal Anti-beta actin antibody (C4) HRP-conjugated antibody | Santa Cruz | sc-47778 HRP |
| Donkey IRDye® 800CW anti-rabbit IgG secondary antibody | LI-COR Biotech | 926-32213 |
| Goat IRDye 680RD anti-mouse IgG secondary antibody | LI-COR Biotech | 926-68070 |

### Methods

#### Cell culture

Cell lines used in this study were acquired from the American Type Culture Collection (ATCC), National Cancer Institute Developmental Therapeutics Program (NCI-DTP), or Takara and cultured in a humidified incubator at 37 °C with 5% $CO_2$. Ramos (ATCC, CRL-1596), 22Rv1 (ATCC, CRL-2505), UACC-257 (NCI-DTP, CVCL_1779), U-2 OS (ATCC, HTB-96), and Hep G2 (ATCC, HB-8065) cell lines were maintained in RPMI-1640 supplemented with 10% FBS, 2 mM GlutaMAX™, 100 U/mL penicillin, and 100 µg/mL streptomycin. HEK293T (ATCC, CRL-3216) and Lenti-X (Takara, 632180) cells were maintained in DMEM supplemented with 10% FBS, 2 mM GlutaMAX™, 100 U/mL penicillin, and 100 µg/mL streptomycin.

#### Gel-ABPP for proteome-wide reactivity

Ramos cells (1 mL, 3 million cells/mL) were treated with alkyne probes for 1 h at 37 ˚C. Cells were harvested by centrifugation at 800 × *g* and then washed 2 × 1 mL with ice-cold Dulbecco's phosphate-buffered saline (DPBS). The pellets were resuspended in 200 µL ice-cold DPBS and lysed by probe sonication (2 × 15 pulses, 10% power output). Samples were normalized to 1 mg/mL in 50 µL with the Pierce™ BCA Protein Assay Kit relative to a bovine serum albumin (BSA) standard according to the manufacturer's protocol. Click reactions[1, 2] were subsequently performed by adding 5 µL 'click' mixture [3 µL of 1.7 mM tris[(1-benzyl-1H-1, 3-triazol-4-yl)methyl]amine (TBTA) in 4/1 *t*-BuOH/DMSO v/v, 1 µL of 50 mM $CuSO_4$ in HPLC $H_2O$, 1 µL of 1.25 mM rhodamine-PEG$_3$-azide in DMSO, 1 µL of freshly prepared 50 mM tris-(2-carboxyethyl)phosphine hydrochloride (TCEP) in HPLC water] for 1 h followed by the addition of 14 µL 4× LDS gel loading buffer containing 5% v/v 2-mercaptoethanol (BME). Samples were denatured at 95 ˚C, resolved at 275 V for 240 min on a 10% tris-glycine gel, and the in-gel fluorescence was measured with a ChemiDoc MP imaging system using the rhodamine filter set. Gels were then stained with Coomassie Instant Blue™ for 10 min at room temperature, destained for 5 min in water, and visualized with a ChemiDoc MP imaging system using the Coomassie filter. Images were processed using Image Lab software (version 6.1.0).

#### Protein-directed ABPP in pooled cells

***In situ* compound treatment and sample processing.** Ramos, 22Rv1, UACC-257, U-2 OS, and Hep G2 cells were seeded and cultured separately under standard conditions. On the day of treatment, adherent cell lines were detached with trypsin-EDTA (0.25%) at 37 ˚C for 5 min and quenched with complete media before pelleting at 800 × *g* for 5 min and resuspending in complete media. Suspension cell lines were pelleted at 800 × *g* for 5 min and resuspended in complete media. Cell lines were then counted on a TC20 automated cell counter with trypan blue (1:1 v/v) and pooled in the appropriate ratio to yield equal protein content per cell line, as measured by Pierce™ BCA Protein Assay Kit relative to a BSA standard. For 1 mg protein per channel: Ramos: 3.75 million/channel; 22Rv1: 1.55 million cells/channel; Hep G2: 0.9 million/channel; UACC-257: 0.54 million/channel; U-2 OS: 1.05 million/channel. Pooled cells were treated as a 10 mL suspension with DMSO or 20 µM non-alkyne competitor stereoprobes (0.1% DMSO v/v final) for 2 h with mixing every 15 min. The cells were then treated with 5 µM stereochemically matched alkyne probes (0.1% DMSO v/v final) for 1 h with mixing every 15 min. The cells were washed 3 × 10 mL with ice-cold DPBS by centrifuging at 800 × *g* for 5 min, transferred into 1.5 mL tubes, and stored at −80 °C. Protein-directed ABPP (i.e., cell lysis, normalization, copper-catalyzed 'click' chemistry-based ligation to biotin-PEG$_4$-azide, streptavidin enrichment, TMTpro16 tag-based multiplexing, Sep-Pak desalting, preparatory HPLC-based high-pH fractionization and LC-MS analysis) was carried out as previously reported[3-5] with the following minor modifications. Pierce™ High Capacity Streptavidin Agarose beads were used to capture liganded proteins for 2 h at room temperature with rotation and, after washing, enriched proteins were digested for 16 h with 75 µL of trypsin mix (200 mM EPPS pH 8.0 supplemented with 1 M urea, 1 mM $CaCl_2$ and 20 µg/mL trypsin) at 37 ˚C with shaking. The beads were pelleted at 5,000 × *g* for 2 min at room temperature, the supernatant was collected, diluted with 25 µL of acetonitrile (25%

v/v final), and labeled with 5 μL of a 20 mg/mL solution of the corresponding TMTpro16 plex tag in anhydrous acetonitrile for 1 h at room temperature with vortexing every 30 min.

**TMT liquid chromatography-mass-spectrometry (LC-MS) analysis.** All LC-MS analysis and data analysis were performed as described previously.[3-6] Briefly, samples were run on an Orbitrap Eclipse Tribrid Mass Spectrometer (Thermo Scientific) or an Orbitrap Fusion Tribrid Mass Spectrometer (Thermo Scientific) coupled to an UltiMate 3000 Series Rapid Separation LC system and autosampler (Thermo Scientific Dionex).

**Data processing.** Raw files were uploaded to the Integrated Proteomics Pipeline (IP2, version 6.0.2) available at http://ip2.scripps.edu/ip2/mainMenu.html, and MS2 and MS3 files were extracted from the raw files using RAW Converter (version 1.1.0.22, available at http://fields.scripps.edu/rawconv/) and searched using the ProLuCID algorithm using a reverse concatenated, non-redundant variant of the Human UniProt database (release 2016-07). Cysteine residues were searched with a static modification for carboxyamidomethylation (+57.02146 Da) and methionine residues were searched with a differential modification for oxidation (+15.9949 Da). Enrichment ratios (probe versus probe) were calculated for each peptide–spectra match by dividing each TMT reporter ion intensity by the sum intensity for all the channels. Peptide–spectra matches were then grouped based on protein ID and (excluding peptides with summed reporter ion intensities less than 10,000, coefficient of variation of greater than 0.5, and less than 2 distinct peptides). Replicate channels were grouped across each experiment and average values were computed for each protein. A variability metric was also computed across replicate channels, which equaled the ratio of the standard deviation to the mean for channels corresponding to the highest enriching probe and was expressed as a percentage. A protein was considered stereoselectively enriched if it was (1) quantified in two independent experiments, (2) the average enrichment by the alkyne probe was greater than 3-fold that of its enantiomer and (3) if the variability corresponding to the alkyne probe with the highest enrichment was less than or equal to 30%. A protein was considered stereoselectively liganded if enrichment was greater than 2-fold the enrichment observed following treatment with a stereochemically matched, non-alkyne competitor (i.e. 50% blockade of enrichment in competitor-treated channels as compared to DMSO-pretreated channels).

**Cross-competition ABPP in pooled cells**

***In situ* compound treatment and sample processing.** Pooled, suspended cells were prepared to the "Protein-directed MS-ABPP in pooled cells" protocol and then treated with DMSO, 20 μM non-alkyne acrylamide competitor, or 20 μM non-alkyne butynamide competitor (0.1% DMSO v/v) for 2 h at 37°C with mixing every 15 min. The cells were then treated with 5 μM stereochemically matched alkyne probes (acrylamide or butynamide) for 1 h at 37°C with mixing every 15 min. Subsequent collection and sample processing were performed as described in "Protein-directed ABPP in pooled cells."

**Data processing.** Enrichment ratios and competitions were calculated using the same workflow as in "Protein-directed ABPP in pooled cells." A protein was considered butynamide-preferring if it showed greater than 2-fold blockade of enrichment by the parent butynamide that was also 1.5-fold greater blockade than that of the stereo-matched parent acrylamide; whereas a protein was considered acrylamide-preferring it showed greater than 2-fold blockade of enrichment by the parent acrylamide that was also 1.5-fold greater blockade than that of the stereo-matched parent butynamide. The shared targets show greater than 2-fold blockade of enrichment by both the parent butynamide and acrylamide. Inconclusive targets are those with less than 50% blockade by either parent acrylamide or butynamide in cross-competition ABPP, or showing conflicting blockade values when enriched by acrylamide or butynamide alkynes (e.g., acrylamide-preferring when enriched with acrylamide alkyne while butynamide-preferring when enriched with butynamide alkyne).

**Cysteine-directed ABPP in pooled cells**

***In situ* treatment and sample processing.** Pooled, suspended cells were prepared to the "Protein-directed MS-ABPP in pooled cells" protocol and then treated with DMSO or 20 μM tryptoline butynamide stereoprobes for 3 h at 37 °C in suspension state with mixing every 15 min. Subsequent sample collection was performed as described in "Protein-directed ABPP in pooled cells." Cysteine-directed ABPP was carried out as previously reported.[3-6]

**Data processing.** Raw files were uploaded to the Integrated Proteomics Pipeline (IP2, version 6.0.2) available at http://ip2.scripps.edu/ip2/mainMenu.html, and MS2 and MS3 files were extracted from the raw files using RAW Converter (version 1.1.0.22, available at http://fields.scripps.edu/rawconv/) and searched using the ProLuCID algorithm using a reverse concatenated, non-redundant variant of the Human UniProt database (release 2016-07). Cysteine residues were searched with a static modification for carboxyamidomethylation (+57.02146 Da). A dynamic modification for IA-DTB labeling (+398.25292 Da) was included with a maximum number of two differential modifications per peptide. Cysteine engagement ratios (DMSO versus compound) were calculated for each peptide–spectra match by dividing each TMT reporter ion intensity by the average intensity for the DMSO channels. Peptide–spectra matches were then grouped based on protein ID and residue number (for example, AGPS_C190), excluding peptides with summed reporter ion intensities for the DMSO channels of less than 10,000 and a coefficient of variation for DMSO channels of greater than 0.5. Replicate channels were grouped across each experiment, and average values were computed for each cysteine. A variability metric was also computed across replicate channels, which equaled the ratio of the median absolute deviation to the average and was expressed as a percentage. A cysteine site was considered enantioselectively liganded if the variability corresponding to the probe leading to the highest blockade of IA-DTB was less than or equal to 20%, and at least one of the following additional criteria were met: (1) the average IA-DTB blockade by the probe was greater than 50% and greater than 1.5-fold of its enantiomer, and protein was stereoselectively liganded by protein-directed ABPP, or (2) IA-DTB blockade by the probe was greater than 66.7% and greater than 2.5-fold that of its enantiomer, and the same probe led to less than 25% IA-DTB blockade of at least one other cysteine in the same protein. Tryptoline acrylamide data were from previously published data and processed in the same manner.

**Targeted Lys-C digested ABPP**

***In vitro* treatment and sample processing.** HNT-34 (DSMZ, ACC 600) cells were lysed and normalized to 2.5 mg/mL. 500 µg total protein was titrated with varying concentrations of WX-02-621 or WX-02-622 (500–0.8 µM, 5-fold dilutions in duplicates) for 1 h at room temperature. Compound treated lysates were then incubated with DBIA probe for 1 h at room temperature. Samples were digested with LysC and processed with standard protocol described in "Cysteine-directed ABPP." Targeted data acquisition of the HELLS_364 peptide was performed via parallel reaction monitoring (PRM) on an Exploris Orbitrap 120, as previously described.[7, 8] Briefly, precursor ions corresponding to an IA-DTB-modified HELLS_364 peptide (amino acids 364–371, +3 charge state, m/z 452.28) were isolated and fragmented by high energy collision-induced dissociation and fragments were detected in the Orbitrap at 17,500 resolution. The resulting PRM data were analyzed using Skyline and quantification was performed by summing the peak areas corresponding to six fragment ions from each peptide.

**Data processing.** Raw AUCs were normalized using RTS peptides (#2–11) and target engagement was calculated relative to DMSO controls ($n$ = 16).

**IP-MS**

***In situ* treatment and sample processing.** Cells stably expressing FLAG epitope-tagged protein of interest (HEK293T ACTMAP WT or C132A) were seeded at 5 million cells per 10 cm plate the day prior to treatment. The media was removed and replaced with fresh media containing DMSO or 10 µM stereoprobes (0.1% DMSO v/v) for 3 h. The cells were detached with trypsin-EDTA (0.25%), quenched with complete DMEM media, collected by centrifugation at 500 × $g$ for 5 min, and washed 3 × 10 mL with ice-cold DPBS. Cell pellets were stored at −80 ˚C, thawed in 1 mL of IP lysis buffer [50 mM EPPS pH 8.0 supplemented with 150 mM NaCl, 1% Triton™ X-100 v/v, 10% glycerol v/v, and one EDTA-free cOmplete protease inhibitor per 10 mL], and lysed by rotating for 1 h at 4 °C. The lysate was clarified by centrifugation at 16,000 × $g$ for 5 min at 4 ˚C and the supernatant was normalized to 1–3 mg in 1 mL with the Pierce™ BCA Protein Assay Kit relative to a BSA standard. Each sample was treated with 40 µL prewashed anti-FLAG magnetic beads for 3 h at 4 ˚C with rotation. The samples were washed 3 × 1 mL with IP wash buffer (25 mM EPPS pH 8.0 supplemented with 150 mM NaCl, 0.2% Triton™ X-100 v/v) and 1 × 1 mL with 50 mM EPPS pH 8.0 using a DynaMag-2 magnetic rack. Enriched proteins were eluted in 40 µL 50 mM EPPS pH 8.0 supplemented with 8 M urea at 65 ˚C for 10 min. The supernatant was collected and reduced by adding 2 µL 200 mM DTT in HPLC water for 15 min at 65 ˚C and alkylated by adding 2 µL 400 mM iodoacetamide in HPLC water for 30 min at 37 ˚C. Samples were diluted to 2 M urea by adding 115 µL 50 mM EPPS pH 8.0 and digested with 4 µL trypsin mix (50mM EPPS supplemented with 25 mM $CaCl_2$ and 0.25 µg/µL trypsin) overnight at 37 ˚C with shaking. Acetonitrile (75 µL) was added, and the samples were multiplexed with TMTpro 16plex and desalted as described in "Protein-directed ABPP in pooled cells." The multiplex sample was then high-pH spin column fractionated and pooled into three fraction as previously described.[3-5]

**Data processing.** IP-MS data were processed in the same manner as described in "Protein-directed ABPP pooled cells" except protein signals were normalized to the FLAG epitope-tagged protein pulled down within each treatment group and then to DMSO signals across treatment groups, unless otherwise noted.

**Time-course and Dosage MS-ABPP for ACTMAP liganding.**
10 million 22Rv1 cells were seeded per 10 cm plate 12 h before treatment. Cells were treated with alkynes at 10 µM for 2–12 h (time-course) or 1–20 µM (concentration screen) for 2 h before adding 5 µM alkyne for 1 h. The samples were collected and processed in the same workflow as described in "Protein-directed ABPP in pooled cells."
**Data processing.** Processing and analysis of MS data followed the "Protein-directed ABPP in pooled cells." For time-course MS-ABPP, only proteins that were enantioselectively enriched by WX-02-570 or WX-02-571 in "Protein-directed ABPP in pooled cells" were analyzed and plotted for enrichment changes at 8 h versus 2 h.

**Cloning and mutagenesis**
AGPS plasmids were obtained from GenScript in pcDNA3.1-C-(k) DYK (FLAG). C19orf54/ACTMAP and HELLS plasmids were obtained from Twist Biosciences after codon optimization with C-FLAG in entry clones (pTwist ENTR). Mutagenesis was performed with Q5® Site-Directed Mutagenesis Kit, using primers shown below. Wild-type and cysteine-to-alanine/serine mutants of C-terminally FLAG-tagged HELLS and ACTMAP were prepared by Gateway cloning from pTwist ENTR into lentiviral vector pLX304 (Addgene, 25890) using Gateway™ LR Clonase™ II Enzyme mix. sgRNAs of ACTMAP (sequences reported below) were annealed and cloned into LentiCRISPR-v2 (Addgene, 52961) using a previously reported protocol[9] except BsmBI was from NEB and the digestion was performed according to the manufacturer's protocol.

**Table S3:** Primers used for Q5® Site-Directed Mutagenesis.

| Gene | Mutant | Forward primer (5'– 3') | Reverse primer (5'– 3') |
|---|---|---|---|
| HELLS | C364S | GAACATGAAGAGCCGTCTCATCC | TTAATGCGATGTCCC |
| ACTMAP | C132A | GGGCCCTCAAGCGGGTCTGGTGG | TCTTGAATCAAACTCGGAAG |
| ACTMAP | C119A | GATTCTGTTTGCGGCCGACCTTCCG | CACCTCAGGTCTCCTC |
| ACTMAP | C271A | TGGTACACCCGCGCAACCACCAAG | AGCACTGGGTGGAAG |
| AGPS | C190A | TCATGGTCATGCGCTTCATGAGATATTTTTGC | GCTCTAAATACTCGATCATC |

**Table S4:** sgRNAs cloned into LentiCRISPR-v2 for ACTMAP knockout.

| ACTMAP | sgRNA |
| --- | --- |
| sgRNA1 | ACCCGTATGAGTCTCTCCAG |
| sgRNA2 | GGACGCCACTGGGGGGCGAC |
| sgRNA3 | GAAACTACACAAGGACCGGT |
| sgRNA5 | ACCGGGCCGGCCTTATCAA |

**Generation of ACTMAP stable HEK293T cell lines**

Lentivirus was prepared by seeding 4 million Lenti-X 293T cells in 10 cm dishes in 8 mL media the day before transfection. 3 µL of FuGENE HD was dissolved in 160 µL OPTI-MEM medium followed by protein-encoding viral vector (3.4 µg), lentiviral packaging vector (pCMV-dR8.91, 1.7 µg) and envelope vector (VSV-G, 0.85 µg). The transfection mix was incubated at room temperature for 10 min before adding dropwise to the Lenti-X 293T cells. The media containing virus was collected 72 h post-transfection, filtered through a 0.45-µm syringe filter, diluted 3:1 with Lenti-X Concentrator and incubated at 4 ˚C with rotation for 2 h. The virus was concentrated by centrifugation at 1,500 × *g* at 4 ˚C for 45 min. The viral pellet was resuspended in 10% volume of original volume with fresh DMEM media. For each transduction, 0.5–1 million HEK293T cells were mixed with 100–200 µL of concentrated viral supernatant in 1–2 mL DMEM media supplemented with 8 µg/mL polybrene in a 12-well plate. HEK293T cells were spin-infected at 1,000 × *g* at 30 ˚C for 1–2 h and incubated for 24 h at 37 ˚C. Selection was initiated with 8 µg/mL blasticidin for 1–2 weeks until a negative control had no surviving cells. Successful transductions were confirmed by FLAG immunoblots (Western blotting method is described in following section "Gel-ABPP with recombinant proteins without immunoprecipitation") and reselected with 8 µg/mL blasticidin if thawing from frozen stocks.

**Generation of ACTMAP CRISPR/Cas9 knockout cells**

Viral supernatants were generated by the transduction of cells with LentiCRISPR-v2 carrying sgACTMAP and stable knock-out cells were generated as described in "Generation of ACTMAP stable cell lines." Stably knocked-out ACTMAP cell lines were selected with 1 µg/mL of puromycin for one week until a negative control had no surviving cells. KO populations were confirmed by immunoblot with anti-ACTMAP antibody and reselected with 1 µg/mL of puromycin if thawing from frozen stocks.

**Gel-ABPP with recombinant proteins without immunoprecipitation**

0.5 million HEK293T cells were seeded per 6-well dish the day before transfection. FLAG-epitope tag plasmids (1–2 µg) were incubated at room temperature with polyethylenimine (PEI, 40,000 Da), 6 µg, 3:1 ratio to plasmid) in 200 µL OPTI-MEM medium for 15 min. The mixture was added dropwise to the cells and allowed to incubate for 24 h under standard culture conditions. Cells were treated with DMSO or with competitor probes for 2 h, followed by alkyne probes for another 1 h. The cells were then detached by trypsin-EDTA (0.25%), quenched with complete DMEM media, collected at 500 × *g* for 5 min and washed 2 × 1 mL with ice-cold DPBS. Cell pellets were resuspended in 100 µL of ice-cold IP lysis buffer [50 mM EPPS pH 8.0 supplemented with 150 mM NaCl, 1% Triton™ X-100 v/v, 10% glycerol v/v, and one EDTA-free cOmplete protease inhibitor per 10 mL], lysed by sonication (1 × 15 pulses, 10% power output) and centrifuged at 16,000 × *g* for 5 min. Supernatants were normalized to 1 mg/mL in 50 µL using the Pierce™ BCA Protein Assay Kit relative to a BSA standard and treated with 5 µL of click mix as described in "Gel-ABPP for proteome-wide reactivity." The click reaction was quenched by the addition of 18 µL of 4× LDS gel loading buffer containing 5% BME and boiled at 95 ˚C for 5 min. The samples were resolved on a 12% tris-glycine SDS–PAGE gel at 160 V for 60 min and imaged by in-gel fluorescent scanning using a ChemiDoc MP imaging system using the rhodamine filter set. Proteins were transferred onto nitrocellulose or PVDF membrane using Power Blotter (1.3 A each gel for 6 min), blocked with 5% non-fat milk in TBST for 1 h at room temperature or overnight at 4 °C. Membranes were incubated with anti-FLAG antibody (HRP-conjugate, 1:2,000 in 5% milk in TBST) or the loading controls anti-GAPDH, anti-β-actin , or anti-β-tubulins (HRP-conjugates, 1:5,000 in 5% milk in TBST) for 1 h at room temperature or overnight at 4 ˚C. The membranes were washed three times with TBST for 5 min at room temperature and were developed with enhanced chemiluminescent western blot substrate and fluorescence was measured on a ChemiDoc MP imaging system using the chemiluminescence filter set. Images were processed using Image Lab software (version 6.1.0).

**IP-based Gel-ABPP with recombinant proteins**

HEK293T cells were seeded and transfected with C-terminally FLAG-tagged AGPS, HELLS, or ACTMAP mutants according to "Gel-ABPP with recombinant proteins without immunoprecipitation." 1 million HEK293T cells stably expressing C-terminally FLAG-tagged ACTMAP were seeded per 6-well the day before treatment. The cells were treated and collected as described in "Gel-ABPP with recombinant proteins without immunoprecipitation." Cell pellets were resuspended in 100–200 µL IP lysis buffer [50 mM EPPS pH 8.0 supplemented with 150 mM NaCl, 1% Triton™ X-100 v/v, 10% glycerol v.v, and one tablet EDTA-free cOmplete protease inhibitor per 10 mL], lysed by probe sonication (1 × 15 pulses, 10% power output) and centrifuged at 16,000 × *g* for 5 min. Protein concentrations were normalized in IP lysis buffer to 0.5–1 mg/mL in 200 µL with the Pierce™ BCA Protein Assay Kit relative to a BSA standard. 20 µL was removed for the input and treated with 2 µL click mix from "Gel-ABPP with recombinant proteins without immunoprecipitation") for the 1 h before quenching with 5 µL of 4× LDS gel loading buffer containing 5% BME at 95 ˚C for 10 min. The remaining lysate (180 µL) was mixed with 10 µL of anti-FLAG magnetic beads (prewashed with 2 × 1 mL IP wash buffer and resuspended in the original volume in IP lysis buffer) and incubated at 4 ˚C with rotation for 1–2 h. The samples were washed 2 × 1 mL with ice-cold IP wash buffer (25 mM EPPS pH 8.0 supplemented with 150 mM NaCl, 0.2% Triton™ X-100 v/v) and 1 × 1 mL with ice-cold DPBS. Beads were resuspended in 20 µL DPBS for on-bead click reaction with 3 µL of click mix (same mix as "Gel-ABPP with recombinant proteins without immunoprecipitation") for 1 h at room temperature with vortexing every 15 min. The click reactions were quenched with 5 µL of 4× LDS gel loading buffer containing 5% BME and the enriched proteins were eluted by boiling at 95 ˚C for 5 min. The supernatant was collected with a magnetic stand and the samples were resolved on 12% tris-glycine SDS–PAGE gels at 160 V for 60 min. Input and IP samples were imaged and blotted as described in "Gel-ABPP with recombinant proteins without immunoprecipitation."

**Cell proliferation assays**

5,000 22Rv1 cells were seeded per 96-well in 50 µL RPMI media 24 h before treatment. 50 µL of media containing 2× stereoprobes or DMSO (0.078–20 µM final concentrations, 0.1% DMSO v/v) was added and allowed to continue incubating for 72 h. Cell proliferation was measured by 30 µL CellTiter-Glo (CTG) reagent on CLARIOstar Plate Reader (software version 5.40 R3, firmware version 1.21). Raw values were normalized to DMSO-treated wells and analyzed using GraphPad PRISM (software version 10.4.2).

**Western Blots for ACTMAP functional assays**

3 million 22Rv1 cells or 1 million HEK293T stably expressing WT-ACTMAP or C132A-ACTMAP were seeded per 6-well 12 h before treatment. After changing the media, cells were treated with WX-02-623/624 (1–20 µM 48h for concentration-dependent WB, 10 µM 2–48 h for time-course WB, 10 µM 2–24 h for proteasome inhibitors co-treatment, final 0.1% DMSO v/v) with or without degradation inhibitors (1000× dilution, 2–24 h) and were collected and washed as described in "Gel-ABPP with recombinant proteins without immunoprecipitation." Cell pellets were frozen at – 80 ˚C or directly processed as previously described with minor modifications.[10] Briefly, cells were lysed in 100 µL DPBS, normalized to 2 mg/mL in 50 µL with the Pierce™ BCA Protein Analysis Kit relative to a BSA standard, and denatured by adding an equal volume 4× LDS sample buffer containing 200 mM DTT (2× LDS and 100 mM DTT final). The samples were heated at 95 ˚ C for 5 min before resolving on a 12% tris-glycine SDS-PAGE at 160 V for 60 min. The proteins were then transferred onto nitrocellulose membranes using Power Blotter (1.3 A each gel

for 6 min). The membranes were blocked with 5% non-fat milk in TBST before incubating with primary antibody (anti-C19orf54/ACTMAP, 1:1,000; anti-γ-actin, anti-pan-actin, anti-β-tubulin, and anti-immature β-actin, 1:5,000, in 5% milk in TBST) for 2 h at room temperature or overnight at 4 ˚C. The membranes were washed three times with TBST for 5 min before incubating with the corresponding anti-mouse (680RD) or rabbit (800CW) secondary IRDye antibody (1:5,000 in 5% milk in TBST) for 1 h at room temperature. The membranes were washed three times with TBST for 5 min before imaging with an Odyssey CLx Imager (LI-COR Biotech) and images were processed using Image Studio 1.0.20.

**Glutathione (GSH) reactivity assay**.
This assay was performed as described previously.[3, 11] Briefly, GSH was diluted to a final concentration of 50 µM 100 mM Tris pH 8.8 with 30% v/v acetonitrile. In triplicate, 100 µL of the GSH solution was added to a clear 384-well plate. 5 µL of 10 mM stereoprobe were then added to the GSH solution to achieve a final probe concentration of 500 µM, and the reaction was incubated to the 2 h and 6 h time points at room temperature. 5 µL of 100 mM Ellman's reagent was then added to the plate and the absorbance was read at 440 nm. The concentration of GSH remaining was derived from a standard curve and the observed rate ($k_{obs}$/[I]) was calculated assuming pseudo-first-order reaction kinetics from the following equations: $d[GSH]/dt = -k_{obs} \times [GSH]$, $[GSH]_t = [GSH]_{t0} \times e^{-k_{obs}t}$.

**Small-molecule docking on ACTMAP**
A computationally predicted structure of ACTMAP (AF-Q5BKX5-F1-model_v4), retrieved from the AlphaFold Protein Structure Database,[12-14] was truncated at the N-terminus (aa1-68) due to the low structural organization and low prediction confidence of this protein region (pLDDT < 50). The resulting structure was prepared for calculations (Protein Preparation Workflow). Binding sites were mapped (SiteMap, ≥ 15 points per site, more restrictive definition of hydrophobicity, standard grid). The highest-scoring site (SiteScore 1.072, Dscore 1.004) coincided with the ACTMAP_C132 site and was used as reference to generate a docking grid (Glide Receptor Grid Generation). Ligands (WX-02-623 and WX-02-624) were prepared (LigPrep) and docked on ACTMAP using the previously generated grid (non-covalent, Glide XP, 3 output poses per ligand). Docking output poses were scored (Prime MM-GBSA, VSGB solvation model, OPLS4 force field) and compared based on the scoring output (MM-GBSA dG Bind) as well as the distances from the ACTMAP_C132 sulfur atom to the electrophilic carbon atom in the ligands. All calculations were performed on Schrödinger Maestro version 13.9.138, MMshare version 6.5.138, release 2024-1, platform Darwin-x86_64.

**Protein structure figures**
Figures showing structures of AGPS and ACTMAP were generated in Chimera-X (version 1.10).[15]

**Quantification and data visualization**
Band intensities were quantified using Bio-Rad Image Lab software (version 6.1.0). Data visualization in this paper was performed using GraphPad PRISM software version 10.4.2: MS data, public datasets, and band intensities were plotted with mean ± standard deviation (SD); $IC_{50}$ curves were generated using a variable slope nonlinear regression setting with the top and bottom constraints set to 100% and 0%, respectively. Pie charts were generated by Mircrosoft Excel software version 16.76.

**Data availability**
The mass spectrometry proteomics data have been deposited to the ProteomeXchange Consortium via the PRIDE[16] partner repository with the dataset identifier PXD074619. Previously published datasets[3] relevant to this study are under data identifier PXD042541. Raw proteomic files were searched using the ProLuCID algorithm using a reverse concatenated, non-redundant variant of the Human UniProt database (release 2016-07). Processed proteomic data are provided in Supporting Dataset 1.

### Synthesis and Characterization of Novel Compounds

#### General considerations

All NMR spectra were recorded at 298 K unless otherwise noted. $^1$H NMR spectra were recorded on Bruker Avance III 400, Avance III HD 400, or Avance Neo 400 spectrometers ($^1$H, 400 MHz). $^1$H NMR data are reported as follows: chemical shift (δ), multiplicity (s = singlet, d = doublet, t = triplet, m = multiplet; br = broad, or combinations therein), coupling constants, and integration. Chemical shifts are reported in parts per million (ppm) using the appropriate solvent as reference.[17] Analytical supercritical fluid chromatography (SFC) was performed on a Shimadzu LC system (flow rate: 3 mL/min, back pressure: 100 Bar, column temperature: 35 ºC) equipped with a polydiode array detector. Mass measurements for high-resolution mass spectrometry (HRMS) were performed on a Waters Xevo G2-XS TOF calibrated against sodium formate clusters and using a LeuEnk lockmass. Expected monoisotopic masses were calculated using MassLynx 4.1 and the m/z values for calibrant and lockmass were MassLynx-default values.

#### Synthesis of novel compounds

All novel compounds were synthesized from previously reported intermediates[3] **S1** and **S2** as described below.

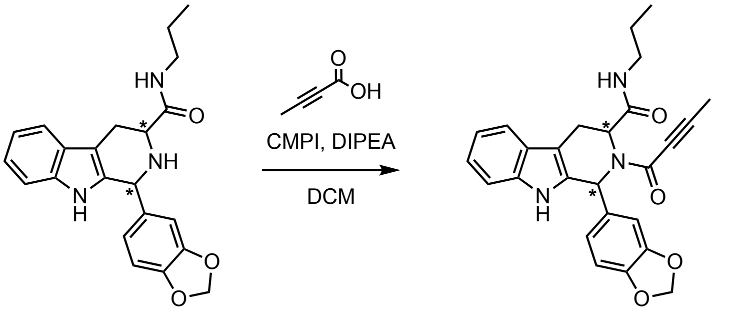


**Scheme S1:** General synthetic scheme for the synthesis of WX-02-621, WX-02-622, WX-02-623, and WX-02-624 from the previously reported intermediates (1*S*,3*S*)-S1, (1*R*,3*R*)-S1, (1*R*,3*S*)-S1, and (1*S*,3*R*)-S1, respectively.

**(1*S*,3*S*)-1-(benzo[d][1,3]dioxol-5-yl)-2-(but-2-ynoyl)-*N*-propyl-2,3,4,9-tetrahydro-1*H*-pyrido[3,4-b]indole-3-carboxamide (WX-02-621)**

To a precooled (0 ˚C) solution of (1*S*,3*S*)-**S1** (80.0 mg, 191 µmol, 1.0 equiv.) and but-2-ynoic acid (32.0 mg, 382 µmol, 2.0 equiv.) in dichloromethane (DCM, 2 mL) were added *N*,*N*-diisopropylethylamine (DIPEA, 74.0 mg, 572 µmol, 3.0 equiv.) and 2-chloro-1-methylpyridinium iodide (CMPI, 73.0 mg, 286 µmol, 1.5 equiv.) at 0 ˚C. The mixture was warmed to 25 ˚C and stirred for 1 h. Upon reaction completion, the mixture was filtered and concentrated under reduced pressure to give a residue, which was purified by prep-HPLC (column: Waters Xbridge 150mm*25mm*5um; mobile phase: [Waters Xbridge 150mm*25mm*5um; mobile phase: A: water (10 mM $NH_4HCO_3$), B: acetonitrile; gradient: 42%–72% B over 9 min) to give **WX-02-621** (24.0 mg, 28% yield) as a white solid.

**$^1$H NMR** (400 MHz, $CD_3OD$) δ 7.60 – 7.47 (m, 1H), 7.31 – 7.23 (m, 1H), 7.15 – 7.07 (m, 1H), 7.07 – 7.01 (m, 1H), 6.91 – 6.66 (m, 4H), 5.94 – 5.86 (m, 2H), 5.64 – 5.49 (m, 1H), 3.69 (dd, *J* = 15.8, 1.5 Hz, 0.6H), 3.51 (dd, *J* = 15.8, 2.6 Hz, 0.4H), 3.05 – 2.91 (m, 1H), 2.91 – 2.77 (m, 1H), 2.54 – 2.42 (m, 0.4H), 2.36 (ddd, *J* = 13.8, 8.4, 6.3 Hz, 0.6H), 2.18 (s, 1.2H), 2.10 (s, 1.8H), 1.30 – 1.07 (m, 2H), 0.80 – 0.66 (m, 3H), 2 exchangeable protons not observed; 6:4 mixture of rotamers.

**HRMS** m/z calc. for $C_{26}H_{26}N_3O_4$ $[M+H]^+$ 444.1923 found 444.1934.

**(1*R*,3*R*)-1-(benzo[d][1,3]dioxol-5-yl)-2-(but-2-ynoyl)-*N*-propyl-2,3,4,9-tetrahydro-1*H*-pyrido[3,4-b]indole-3-carboxamide (WX-02-622)**

To a precooled (0 ˚C) solution of (1*R*,3*R*)-**S1** (75.0 mg, 199 μmol, 1 equiv.) and but-2-ynoic acid (33.4 mg, 397 μmol, 2.0 equiv.) in DCM (2 mL) were added DIPEA (77.0 mg, 596 μmol, 3.0 equiv.) and CMPI (76.2 mg, 298 μmol, 1.5 equiv.). The mixture was stirred at 25 ˚C for 1 h. Upon reaction completion, the reaction mixture was concentrated under reduced pressure to give a residue, which was purified by prep-TLC ($SiO_2$, petroleum ether/EtOAc = 1:1) and prep-HPLC (column: Waters Xbridge 150mm*25mm*5um; mobile phase: A: water (10 mM $NH_4HCO_3$), B: acetonitrile; gradient: 40%–70% B over 9 min) to give **WX-02-622** (37.0 mg, 42% yield) as an off-white solid.

**$^1$H NMR** (400 MHz, $CD_3OD$) δ 7.60 – 7.47 (m, 1H), 7.31 – 7.23 (m, 1H), 7.15 – 7.07 (m, 1H), 7.07 – 7.01 (m, 1H), 6.91 – 6.66 (m, 4H), 5.94 – 5.86 (m, 2H), 5.64 – 5.49 (m, 1H), 3.69 (dd, *J* = 15.8, 1.5 Hz, 0.6H), 3.51 (dd, *J* = 15.8, 2.6 Hz, 0.4H), 3.05 – 2.91 (m, 1H), 2.91 – 2.77 (m, 1H ), 2.54 – 2.42 (m, 0.4H), 2.35 (ddd, *J* = 13.1, 8.4, 6.2 Hz, 0.6H), 2.18 (s, 1.2H), 2.09 (s, 1.8H), 1.30 – 1.07 (m, 2H), 0.78 – 0.69 (m, 3H), 2 exchangeable protons not observed; 6:4 mixture of rotamers.

**HRMS** m/z calc. for $C_{26}H_{26}N_3O_4$ $[M+H]^+$ 444.1923 found 444.1930.

**(1*R*,3*S*)-1-(benzo[d][1,3]dioxol-5-yl)-2-(but-2-ynoyl)-*N*-propyl-2,3,4,9-tetrahydro-1*H*-pyrido[3,4-b]indole-3-carboxamide (WX-02-623)**

To a solution of (1*R*,3*S*)-**S1** (160 mg, 424 μmol, 1.0 equiv.) and but-2-ynoic acid (40.0 mg, 476 μmol, 1.1 equiv.) in DCM (10 mL) were added DIPEA (164 mg, 1.27 mmol, 3.0 equiv.) and CMPI (162 mg, 636 μmol, 1.5 equiv.). The mixture was stirred at 25 ˚C for 1 h. Upon reaction completion, the reaction mixture was diluted with water (30 mL) and extracted with DCM (20 mL × 3). The combined organic layers were dried over anhydrous sodium sulfate, filtered and concentrated under reduced pressure to give a residue, which was purified by prep-TLC ($SiO_2$, EtOAc) and prep-HPLC (column: Waters Xbridge 150mm*25mm*5um; mobile phase: A: water (10 mM $NH_4HCO_3$), B: acetonitrile; gradient: 38%–68% B over 9 min) to give **WX-02-623** (64.0 mg, 34% yield) as a white solid.

**$^1$H NMR** (400 MHz, $CD_3OD$) δ 7.43 – 7.38 (m, 1H), 7.26 (d, *J* = 8.1 Hz, 0.4H), 7.22 (d, *J* = 7.9 Hz, 0.6H), 7.08 – 6.94 (m, 2H), 6.93 – 6.83 (m, 2H), 6.78 (d, *J* = 7.9 Hz, 0.4H), 6.70 (d, *J* = 8.0 Hz, 0.6H), 6.49 (s, 0.4H), 6.18 (s, 0.6H), 5.92 – 5.89 (m, 0.8H), 5.88 – 5.83 (m, 1.2H), 5.54 (dd, *J* = 5.0, 2.8 Hz, 0.6H), 5.14 (dd, *J* = 5.8, 3.8 Hz, 0.4H), 3.59 – 3.37 (m, 2H), 3.07 – 2.98 (m, 2H), 2.02 (s, 1.8H), 1.93 (s, 1.2H), 1.41 – 1.25 (m, 2H), 0.72 (t, *J* = 7.4 Hz, 1.2H), 0.68 (t, *J* = 7.4 Hz, 1.8H), 2 exchangeable protons not observed; 6:4 mixture of rotamers.

**HRMS** m/z calc. for $C_{26}H_{26}N_3O_4$ $[M+H]^+$ 444.1923 found 444.1931.

**(*1S*,3*R*)-1-(benzo[d][1,3]dioxol-5-yl)-2-(but-2-ynoyl)-*N*-propyl-2,3,4,9-tetrahydro-1*H*-pyrido[3,4-b]indole-3-carboxamide (WX-02-624)**

To a solution of (1*S*,3*R*)-**S1** (100 mg, 265 μmol, 1.0 equiv.) and but-2-ynoic acid (44.6 mg, 530 μmol, 2.0 equiv.) in DCM (2 mL) were added DIPEA (103 mg, 795 μmol, 3.0 equiv.) and CMPI (102 mg, 397 μmol, 1.5 equiv.). The mixture was stirred at 25 ˚C for 1 h. Upon reaction completion, the reaction mixture was concentrated under reduced pressure to give a residue, which was purified by prep-TLC ($SiO_2$, petroleum ether/EtOAc = 2:1) and prep-HPLC (column: Waters Xbridge 150mm*25mm*5um; mobile phase: A: water (10 mM $NH_4HCO_3$), B: acetonitrile; gradient: 36%–66% B over 9 min) to give **WX-02-624** (38.0 mg, 85.7 μmol, 32% yield) as an off-white solid.

**$^1$H NMR** (400 MHz, $CD_3OD$) δ 7.43 – 7.38 (m, 1H), 7.26 (dt, *J* = 8.1, 1.0 Hz, 0.4H), 7.22 (dt, *J* = 8.1, 1.0 Hz, 0.6H), 7.08 – 6.94 (m, 2H), 6.93 – 6.83 (m, 2H), 6.78 (d, *J* = 7.9 Hz, 0.4H), 6.69 (d, *J* = 8.0 Hz, 0.6H), 6.49 (s, 0.4H), 6.18 (s, 0.6H), 5.92 – 5.89 (m, 0.8H), 5.85 (dd, *J* = 5.4, 1.2 Hz, 1.2H), 5.54 (dd, *J* = 5.0, 2.8 Hz, 0.6H), 5.14 (dd, *J* = 5.8, 3.8 Hz, 0.4H), 3.57 – 3.36 (m, 2H), 3.07 – 2.98 (m, 2H), 2.02 (s, 1.8H), 1.92 (s, 1.2H), 1.41 – 1.25 (m, 2H), 0.72 (t, *J* = 7.4 Hz, 1.2H), 0.68 (t, *J* = 7.4 Hz, 1.8H), 2 exchangeable protons not observed; 6:4 mixture of rotamers.

**HRMS** m/z calc. for $C_{26}H_{26}N_3O_4$ $[M+H]^+$ 444.1923 found 444.1928.

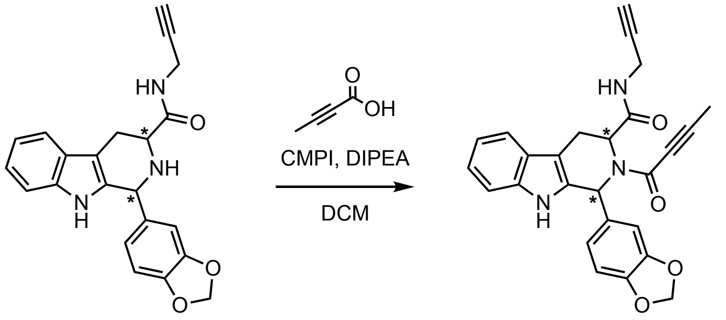


S2 WX-02-568 – WX-02-571

**Scheme S2:** General synthetic scheme for the synthesis of WX-02-568, WX-02-569, WX-02-570, and WX-02-571 from the previously reported intermediates (1*S*,3*S*)-S2, (1*R*,3*R*)-S2, (1*R*,3*S*)-S2, and (1*S*,3*R*)-S2, respectively.

**(1*S*,3*S*)-1-(benzo[d][1,3]dioxol-5-yl)-2-(but-2-ynoyl)-*N*-(prop-2-yn-1-yl)-2,3,4,9-tetrahydro-1*H*-pyrido[3,4-b]indole-3-carboxamide (WX-02-568)**

To a precooled (0 ºC) solution of (1*S*,3*S*)-**S2** (50.0 mg, 134 µmol, 1.0 equiv.), DIPEA (52.0 mg, 201 µmol, 1.5 equiv.) and CMPI (51.0 mg, 201 µmol, 1.5 equiv.) in DCM (1 mL) was added but-2-ynoic acid (23.0 mg, 268 µmol, 2.0 equiv.). The mixture was warmed to 25 ˚C and stirred for 2 h. Upon reaction completion, the mixture was filtered and concentrated to give a residue, which was purified by prep-HPLC (column: Waters Xbridge C18 150mm*50mm*10um; mobile phase: A: water (10 mM $NH_4HCO_3$), B: acetonitrile; gradient: 38%–68% B over 10 min) to obtain **WX-02-568** (8.0 mg, 14% yield) as a white solid.

**$^1$H NMR** (400 MHz, $CD_3OD$) δ 7.58 – 7.50 (m, 1H), 7.32 – 7.23 (m, 1H), 7.16 – 7.08 (m, 1H), 7.07 – 7.00 (m, 1H), 6.92 – 6.61 (m, 4H), 5.95 – 5.90 (m, 2H), 5.64 – 5.56 (m, 1H), 4.59 (s, 1H), 3.74 – 3.49 (m, 2H), 3.22 – 3.08 (m, 1H), 3.04 – 2.90 (m, 1H), 2.50 – 2.44 (m, 1H), 2.19 (s, 1H), 2.10 (s, 2H), 1 exchangeable proton not observed; 2:1 mixture of rotamers.

**HRMS** m/z calc. for $C_{26}H_{22}N_3O_4$ $[M+H]^+$ 440.1610 found 440.1618.

**(1*R*,3*R*)-1-(benzo[d][1,3]dioxol-5-yl)-2-(but-2-ynoyl)-*N*-(prop-2-yn-1-yl)-2,3,4,9-tetrahydro-1*H*-pyrido[3,4-b]indole-3-carboxamide (WX-02-569)**

To a precooled (0 ˚C) solution of (1*R*,3*R*)-**S2** (100.0 mg, 268 µmol, 1.0 equiv.), DIPEA (803 µmol, 140 uL, 3.0 equiv.) and CMPI (103 mg, 402 µmol, 1.5 equiv.) in DCM (2 mL) was added but-2-ynoic acid (45.0 mg, 536 µmol, 2.0 equiv.) at 0 ˚C. The mixture was warmed to 25 ˚C and stirred for 2 h. Upon reaction completion, the reaction mixture was concentrated under reduced pressure to give a residue, which was purified by prep-HPLC (column: Waters Xbridge 150mm*25mm*5um; mobile phase: A: water (10 mM $NH_4HCO_3$), B: acetonitrile; gradient: 35%–65% B over 8 min) to give **WX-02-569** (37.0 mg, 31% yield) as a white solid.

**$^1$H NMR** (400 MHz, $CD_3OD$) δ 7.58 – 7.50 (m, 1H), 7.32 – 7.23 (m, 1H), 7.16 – 7.08 (m, 1H), 7.07 – 7.00 (m, 1H), 6.92 – 6.61 (m, 4H), 5.95 – 5.90 (m, 2H), 5.64 – 5.56 (m, 1H), 4.59 (s, 1H), 3.74 – 3.49 (m, 2H), 3.22 – 3.08 (m, 1H), 3.04 – 2.90 (m, 1H), 2.50 – 2.44 (m, 1H), 2.18 (s, 1H), 2.10 (s, 2H), 1 exchangeable proton not observed; 2:1 mixture of rotamers.

**HRMS** m/z calc. for $C_{26}H_{22}N_3O_4$ $[M+H]^+$ 440.1610 found 440.1613.

**(1*R*,3*S*)-1-(benzo[d][1,3]dioxol-5-yl)-2-(but-2-ynoyl)-*N*-(prop-2-yn-1-yl)-2,3,4,9-tetrahydro-1*H*-pyrido[3,4-b]indole-3-carboxamide (WX-02-570)**

To a precooled (0 ˚C) solution of (1*R*,3*S*)-**S2** (125 mg, 335 µmol, 1.0 equiv.), DIPEA (173 mg, 1.3 mmol, 3.9 equiv.) and CMPI (171 mg, 700 µmol, 2.0 equiv.) in DCM (2 mL) was added but-2-ynoic acid (56 mg, 700 µmol, 2.0 equiv.). The mixture was warmed to 25 ˚C and stirred for 2 h. Upon reaction completion, the

mixture was filtered and concentrated to give a residue, which was purified by prep-HPLC (column: Waters Xbridge C18 150mm*50mm*10um; mobile phase: A: water (10 mM $NH_4HCO_3$), B: acetonitrile; gradient: 34%–64% B over 10 min) to give **WX-02-570** (20.0 mg, 14% yield) as a white solid.
**$^1$H NMR** (400 MHz, $CD_3OD$) δ 7.46 – 7.37 (m, 1H), 7.26 (d, *J* = 8.0 Hz, 0.4H), 7.22 (d, *J* = 7.8 Hz, 0.6H), 7.09 – 6.94 (m, 2H), 6.93 – 6.82 (m, 2H), 6.78 (d, *J* = 8.0 Hz, 0.4H), 6.70 (d, *J* = 8.0 Hz, 0.6H), 6.50 (s, 0.4H), 6.19 (s, 0.6H), 5.93 – 5.89 (m, 0.8H), 5.87 – 5.83 (m, 1.2H), 5.53 (t, *J* = 4.0 Hz, 0.6H), 5.09 (dd, *J* = 5.8, 4.3 Hz, 0.4H), 4.59 (s, 1H), 3.94 – 3.73 (m, 2H), 3.49 (d, *J* = 4.1 Hz, 1.2H), 3.42 – 3.23 (m, 0.8H + residual NMR solvent), 2.48 (t, *J* = 2.5 Hz, 0.6H), 2.44 (t, *J* = 2.6 Hz, 0.4H), 2.03 (s, 1.8H), 1.94 (s, 1.2H), 1 exchangeable proton not observed; 6:4 mixture of rotamers.
**HRMS** m/z calc. for $C_{26}H_{22}N_3O_4$ $[M+H]^+$ 440.1610 found 440.1612.

**(1*S*,3*R*)-1-(benzo[d][1,3]dioxol-5-yl)-2-(but-2-ynoyl)-*N*-(prop-2-yn-1-yl)-2,3,4,9-tetrahydro-1*H*-pyrido[3,4-b]indole-3-carboxamide (WX-02-571)**
To a precooled (0 ˚C) solution of (1*S*,3*R*)-**S2** (70.0 mg, 187 µmol, 1.0 equiv.), DIPEA (72.7 mg, 0.56 mmol, 3.0 equiv.) and CMPI (71.8 mg, 281 µmol, 1.5 equiv.) in DCM (2 mL) was added but-2-ynoic acid (31.5 mg, 375 µmol, 2.0 equiv.). The mixture was warmed to 25 ˚C and stirred for 2 h. Upon reaction completion, the mixture was filtered and concentrated to give a residue, which was purified by prep-HPLC (column: Waters Xbridge 150mm*25mm* 5um; mobile phase: A: water (10 mM $NH_4HCO_3$), B: acetonitrile; gradient: 33%–63% B over 8 min) to give **WX-02-571** (20.0 mg, 24% yield) as a white solid.
**$^1$H NMR** (400 MHz, $CD_3OD$) δ 7.46 – 7.37 (m, 1H), 7.26 (d, *J* = 8.1 Hz, 0.4H), 7.22 (d, *J* = 8.0 Hz, 0.6H), 7.09 – 6.94 (m, 2H), 6.93 – 6.82 (m, 2H), 6.77 (d, *J* = 8.0 Hz, 0.4H), 6.69 (d, *J* = 8.0 Hz, 0.6H), 6.50 (s, 0.4H), 6.20 (s, 0.6H), 5.93 – 5.89 (m, 0.8H), 5.87 – 5.82 (m, 1.2H), 5.53 (t, *J* = 4.0 Hz, 0.6H), 5.09 (t, *J* = 5.0 Hz, 0.4H), 4.59 (s, 1H, partially exchanged), 3.94 – 3.73 (m, 2H), 3.49 (d, *J* = 4.1 Hz, 1.2H), 3.42 – 3.23 (m, 0.8H + residual NMR solvent), 2.48 (t, *J* = 2.6 Hz, 0.6H), 2.44 (t, *J* = 2.5 Hz, 0.4H), 2.03 (s, 1.8H), 1.93 (s, 1.2H), 1 exchangeable proton not observed; 6:4 mixture of rotamers.
**HRMS** m/z calc. for $C_{26}H_{22}N_3O_4$ $[M+H]^+$ 440.1610 found 440.1615.

**Analytical data: NMR spectra**

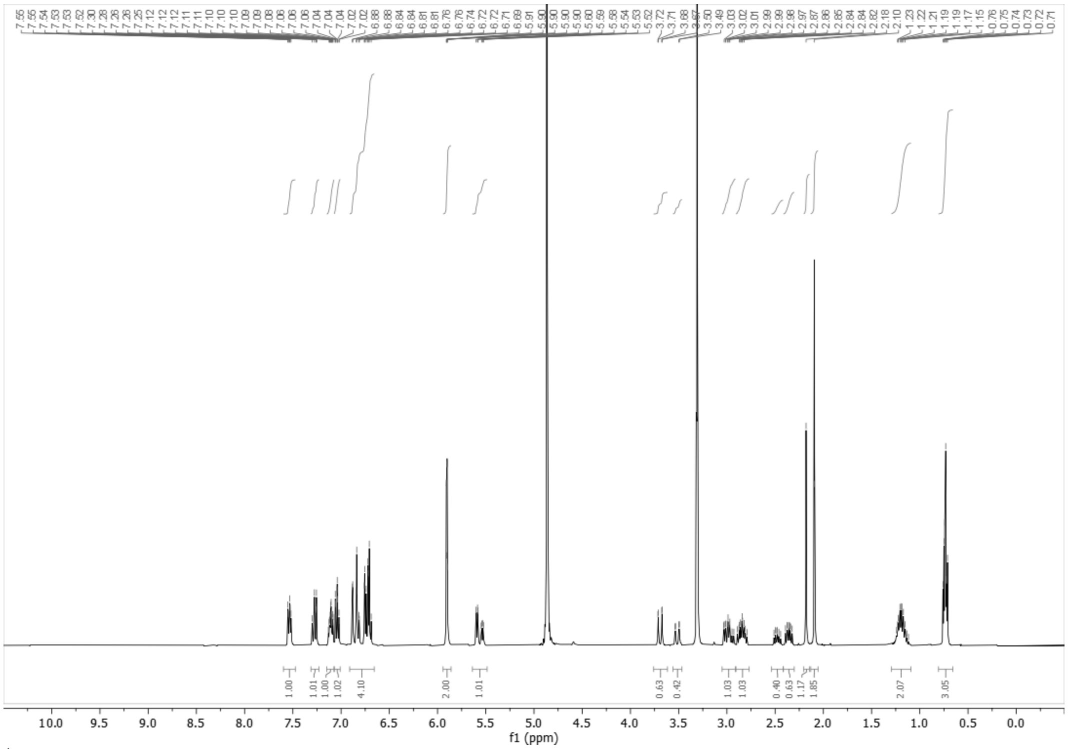

$^1$H NMR spectrum of WX-02-621 (400 MHz, $CD_3OD$)

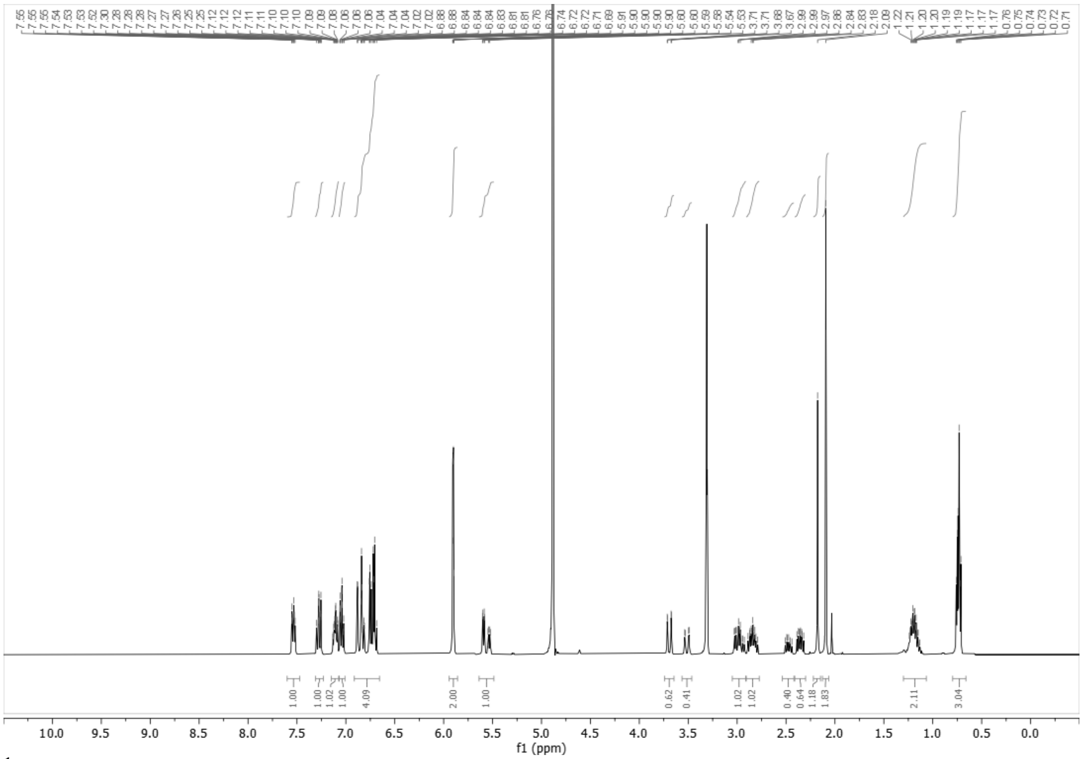

$^1$H NMR spectrum of WX-02-622 (400 MHz, $CD_3OD$)

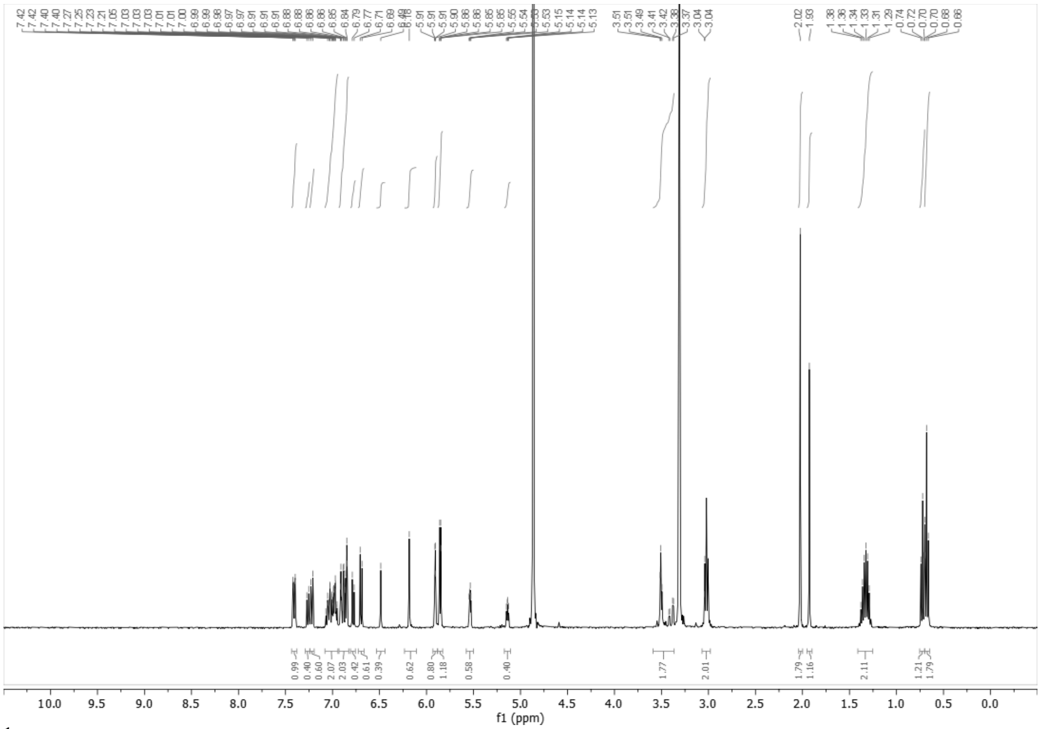

[1]H NMR spectrum of WX-02-623 (400 MHz, $CD_3OD$)

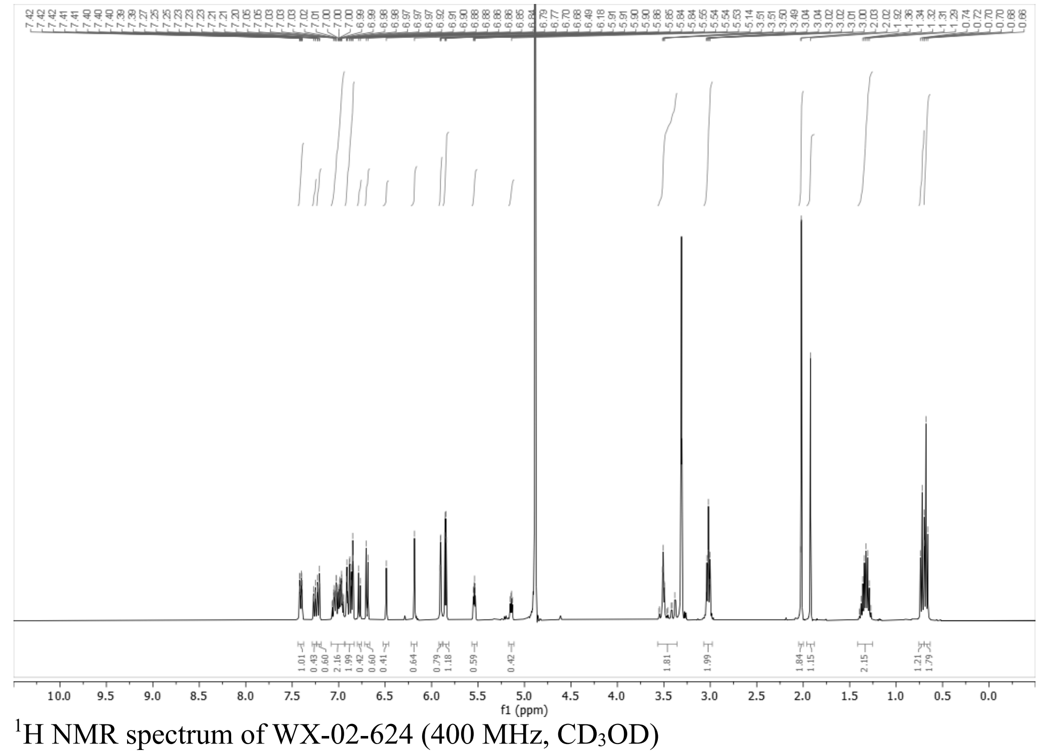

[1]H NMR spectrum of WX-02-624 (400 MHz, $CD_3OD$)

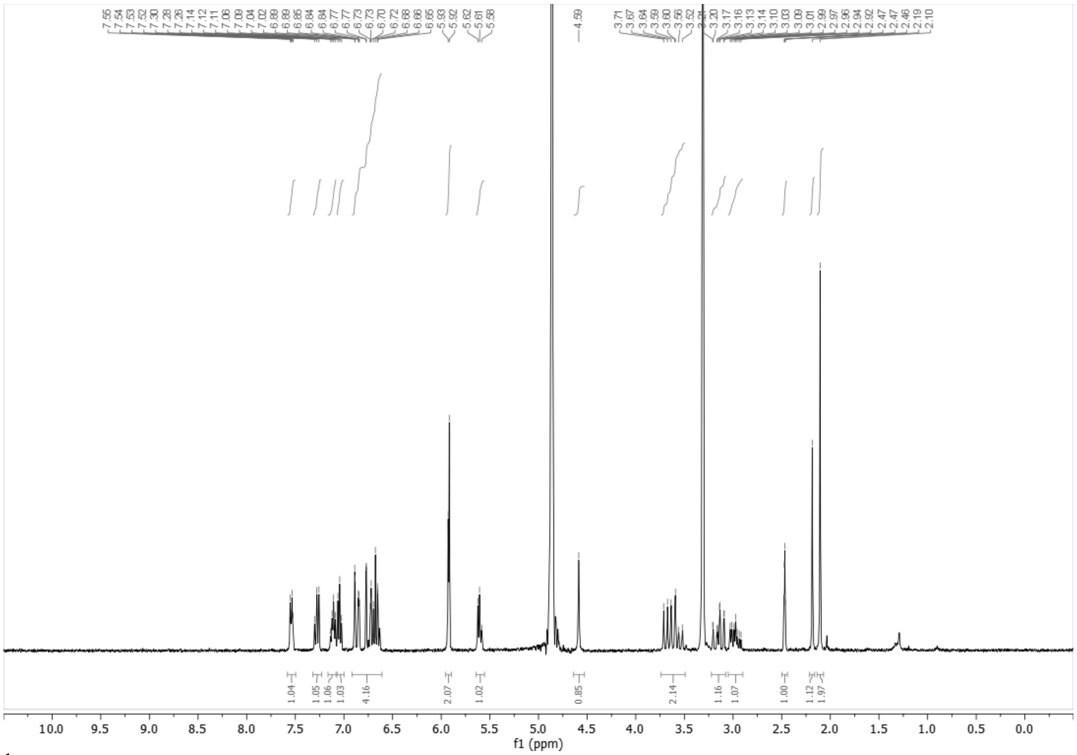


$^1$H NMR spectrum of WX-02-568 (400 MHz, $CD_3OD$)

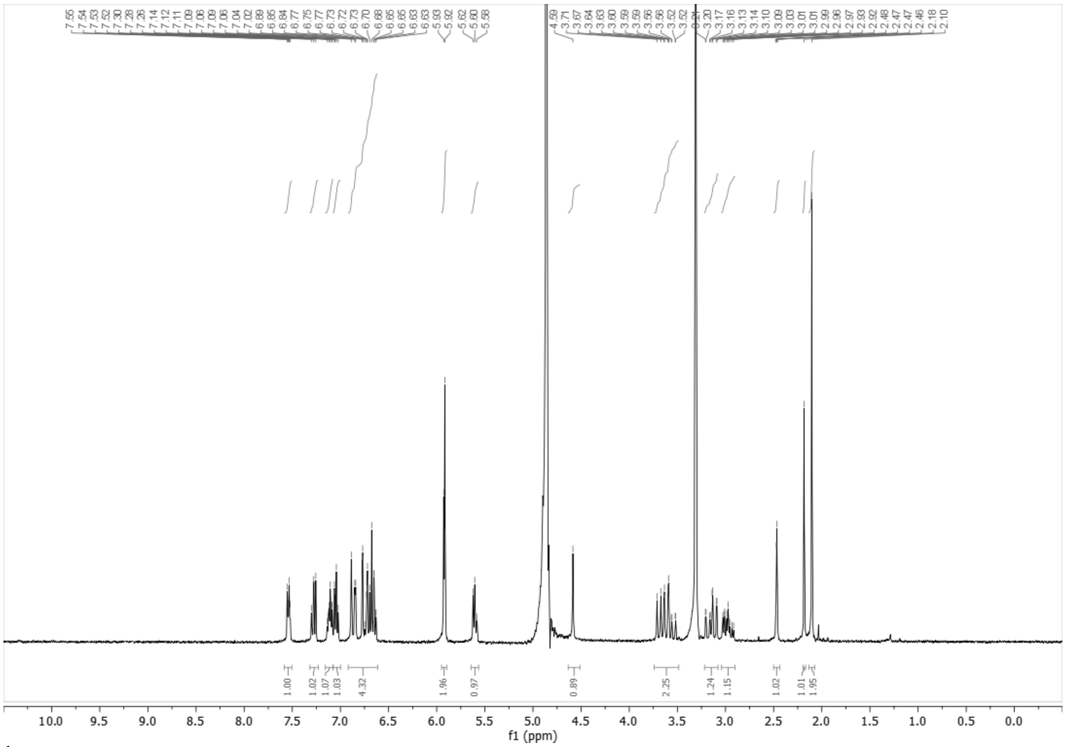


$^1$H NMR spectrum of WX-02-569 (400 MHz, $CD_3OD$)

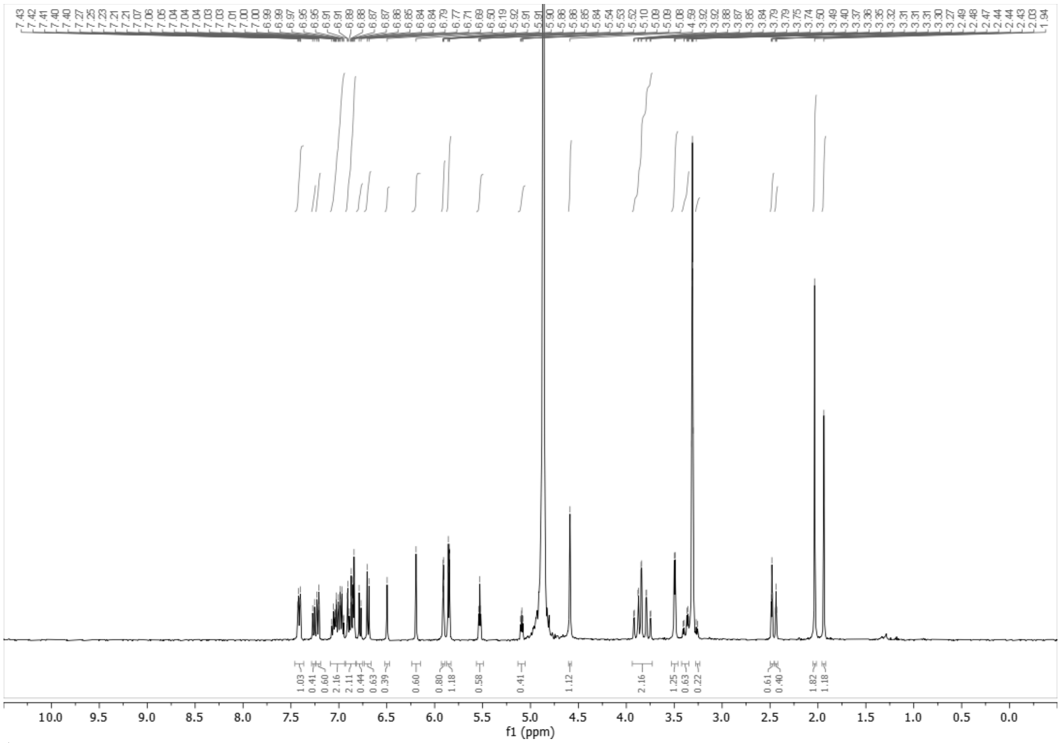

[1]H NMR spectrum of WX-02-570 (400 MHz, $CD_3OD$)

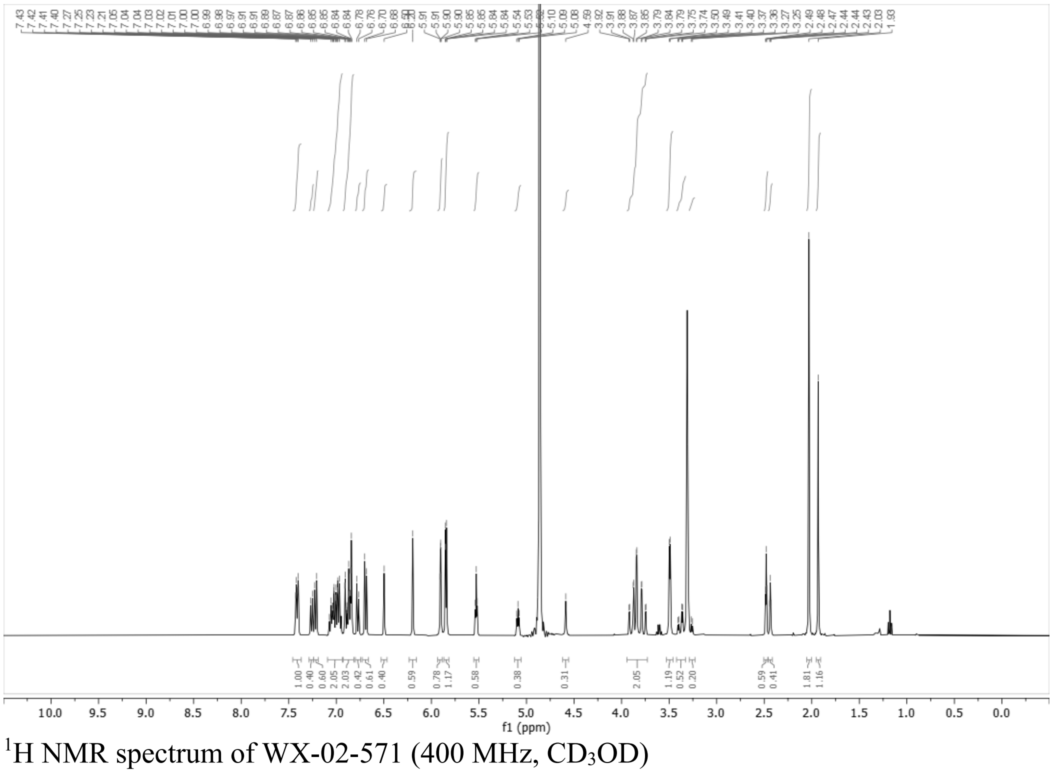

[1]H NMR spectrum of WX-02-571 (400 MHz, $CD_3OD$)

**Analytical data: SFC**

WX-02-621

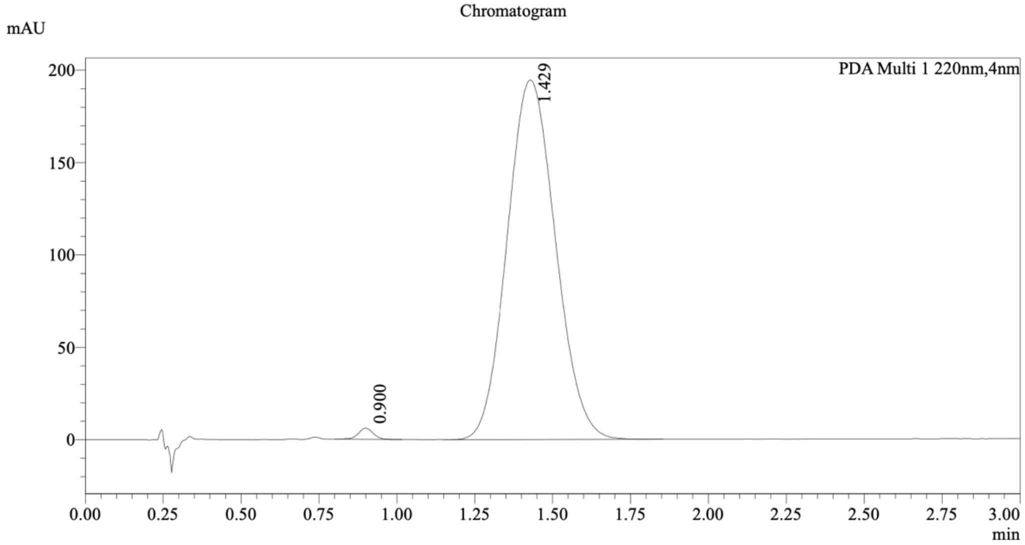


1 PDA Multi 1 / 220nm,4nm

Integration Result

Peak Table

PDA Ch1 220nm

| Peak# | Ret. Time | Height | Height% | Resolution(USP) | Area | Area% |
|---|---|---|---|---|---|---|
| 1 | 0.900 | 6008 | 2.996 | -- | 19972 | 0.940 |
| 2 | 1.429 | 194519 | 97.004 | 2.819 | 2104453 | 99.060 |

WX-02-622

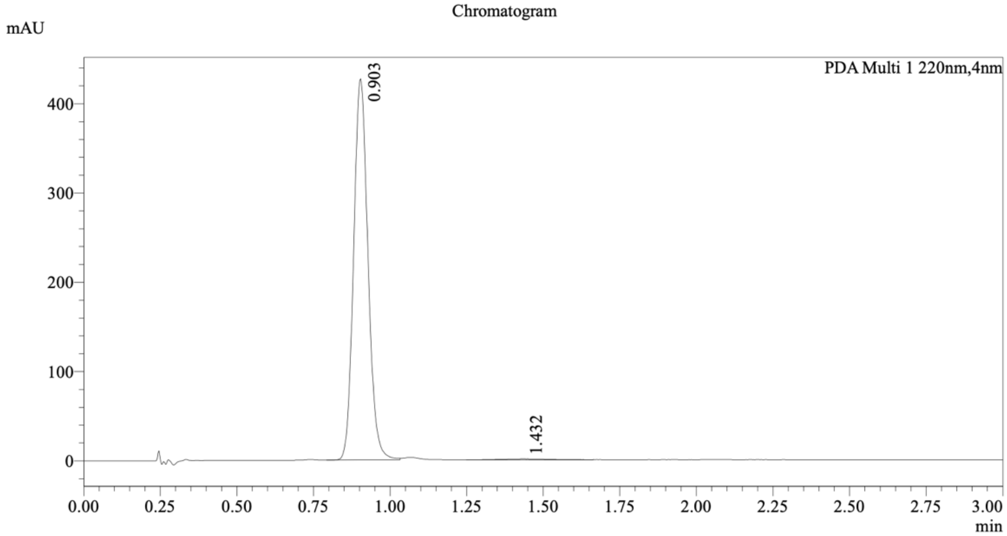


1 PDA Multi 1 / 220nm,4nm

Integration Result

Peak Table

PDA Ch1 220nm

| Peak# | Ret. Time | Height | Height% | Resolution(USP) | Area | Area% |
|---|---|---|---|---|---|---|
| 1 | 0.903 | 423239 | 99.778 | -- | 1389675 | 99.312 |
| 2 | 1.432 | 943 | 0.222 | 2.926 | 9624 | 0.688 |

Mixture of WX-02-621 and WX-02-622

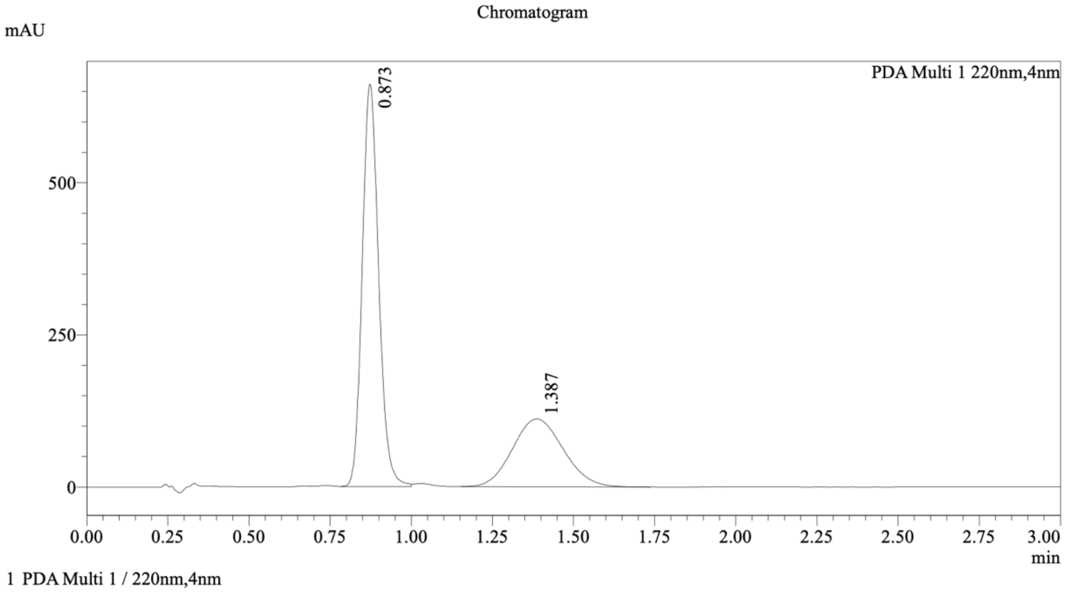


Integration Result

Peak Table

PDA Ch1 220nm

| Peak# | Ret. Time | Height | Height% | Resolution(USP) | Area | Area% |
|---|---|---|---|---|---|---|
| 1 | 0.873 | 651611 | 85.416 | -- | 2330388 | 65.743 |
| 2 | 1.387 | 111260 | 14.584 | 2.670 | 1214286 | 34.257 |

Method Details

Column: Chiralpak IG-3 50×4.6mm I.D., 3 µm
Mobile phase: A: $CO_2$, B: EtOH (0.05% diethylamine)
Elution: 40% B

WX-02-623

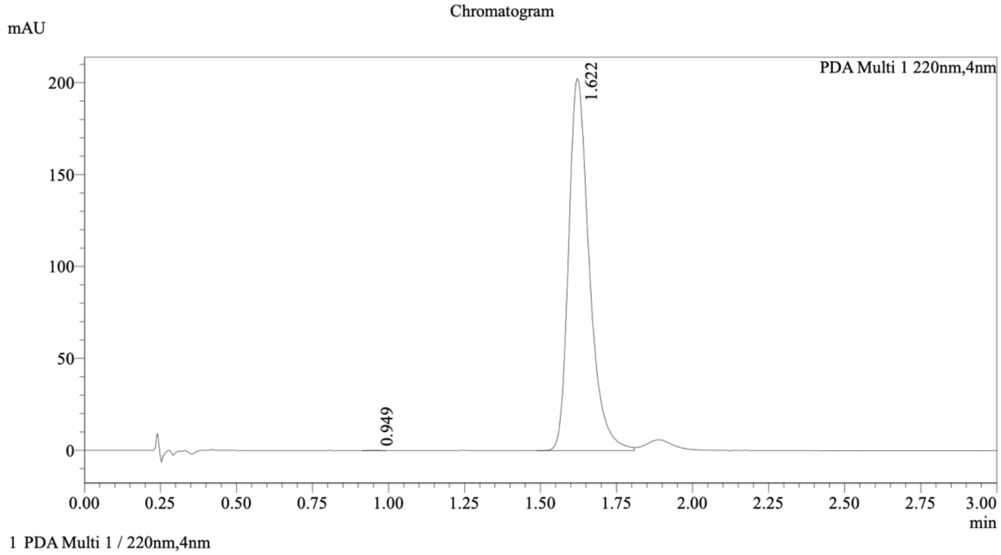


Integration Result

Peak Table

PDA Ch1 220nm

| Peak# | Ret. Time | Height | Height% | Resolution(USP) | Area | Area% |
|---|---|---|---|---|---|---|
| 1 | 0.949 | 126 | 0.062 | -- | 266 | 0.028 |
| 2 | 1.622 | 201816 | 99.938 | 7.293 | 956263 | 99.972 |

WX-02-624

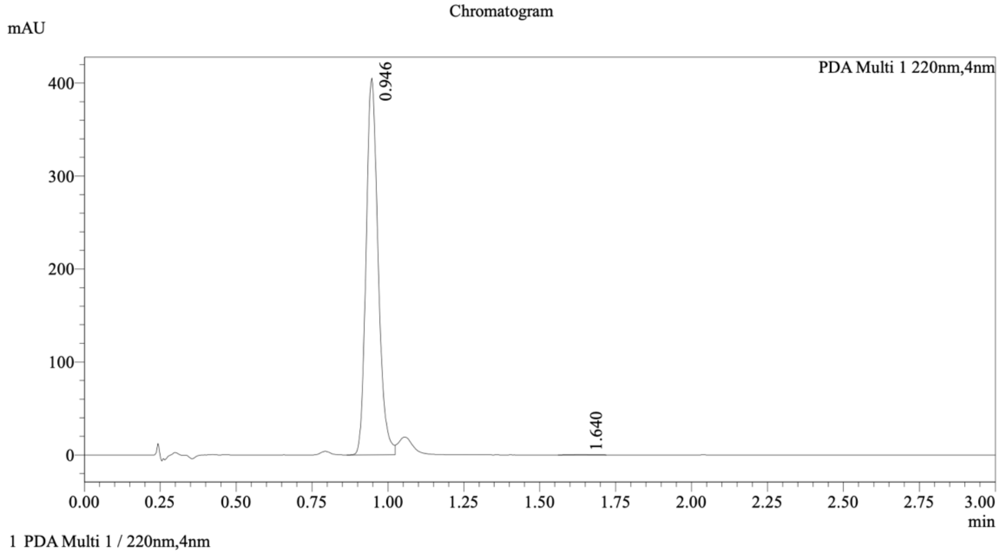


Integration Result

Peak Table

PDA Ch1 220nm

| Peak# | Ret. Time | Height | Height% | Resolution(USP) | Area | Area% |
|---|---|---|---|---|---|---|
| 1 | 0.946 | 395675 | 99.906 | -- | 1114322 | 99.855 |
| 2 | 1.640 | 372 | 0.094 | 7.197 | 1613 | 0.145 |

Mixture of WX-02-623 and WX-02-624

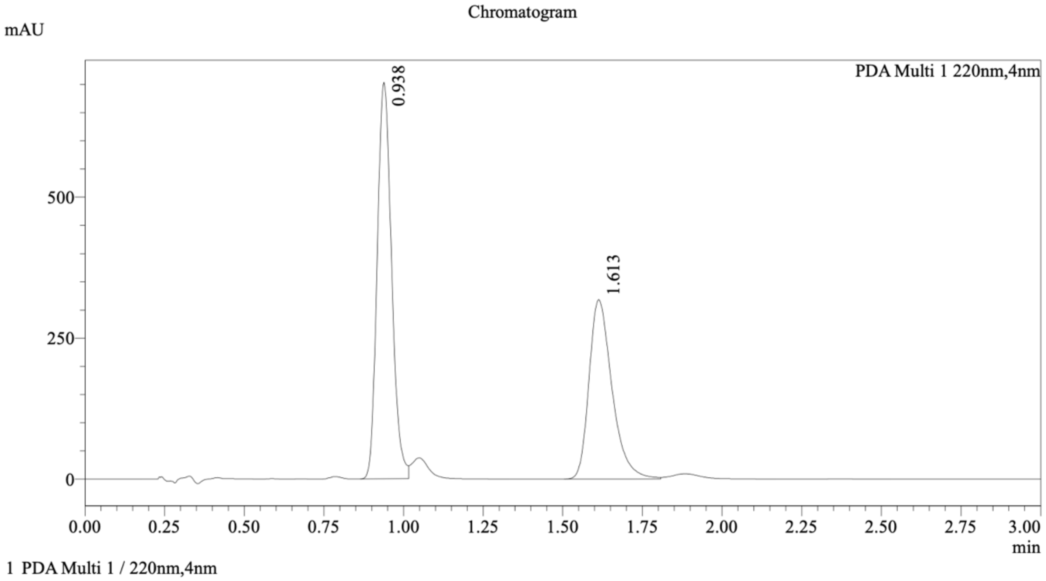


Integration Result

Peak Table

PDA Ch1 220nm

| Peak# | Ret. Time | Height | Height% | Resolution(USP) | Area | Area% |
|---|---|---|---|---|---|---|
| 1 | 0.938 | 691948 | 68.549 | -- | 2209080 | 58.013 |
| 2 | 1.613 | 317479 | 31.451 | 6.279 | 1598812 | 41.987 |

Method Details

Column: (S,S)Whelk-O1 50×4.6mm I.D., 3.5 µm
Mobile phase: A: $CO_2$, B: EtOH (0.05% diethylamine)
Elution: 40% B

WX-02-568

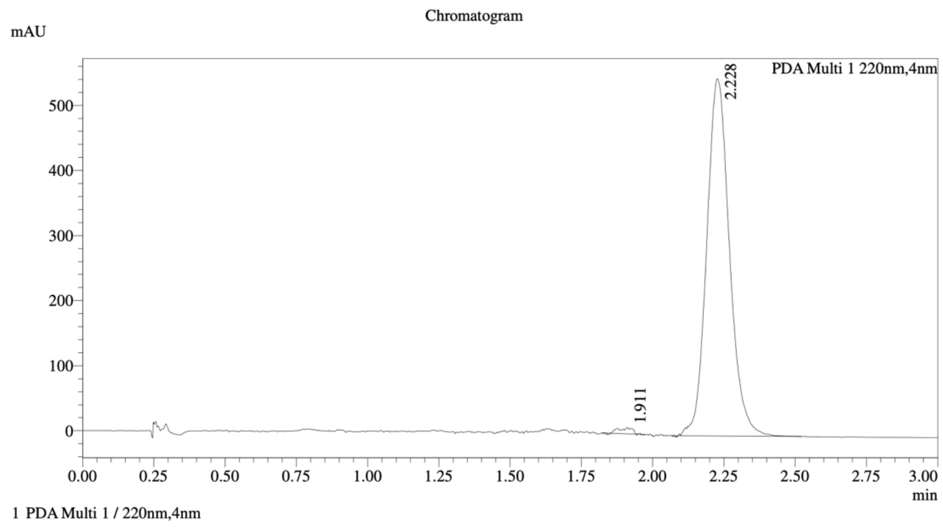


Integration Result

Peak Table

PDA Ch1 220nm

| Peak# | Ret. Time | Height | Height% | Resolution(USP) | Area | Area% |
|---|---|---|---|---|---|---|
| 1 | 1.911 | 9367 | 1.683 | -- | 30758 | 0.997 |
| 2 | 2.228 | 547359 | 98.317 | 3.140 | 3054527 | 99.003 |

WX-02-569

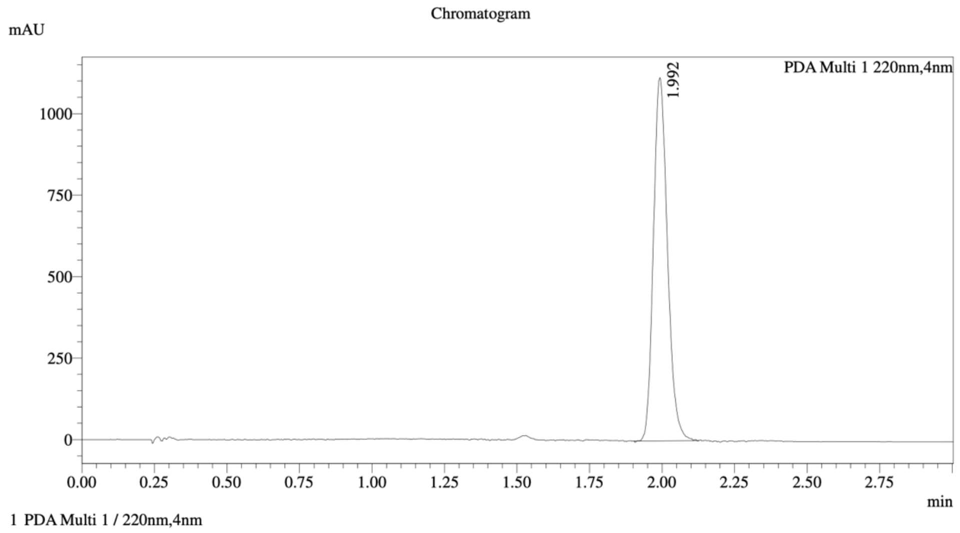


Integration Result

Peak Table

PDA Ch1 220nm

| Peak# | Ret. Time | Height | Height% | Resolution(USP) | Area | Area% |
|---|---|---|---|---|---|---|
| 1 | 1.992 | 1098053 | 100.000 | -- | 3755129 | 100.000 |

Mixture of WX-02-568 and WX-02-569

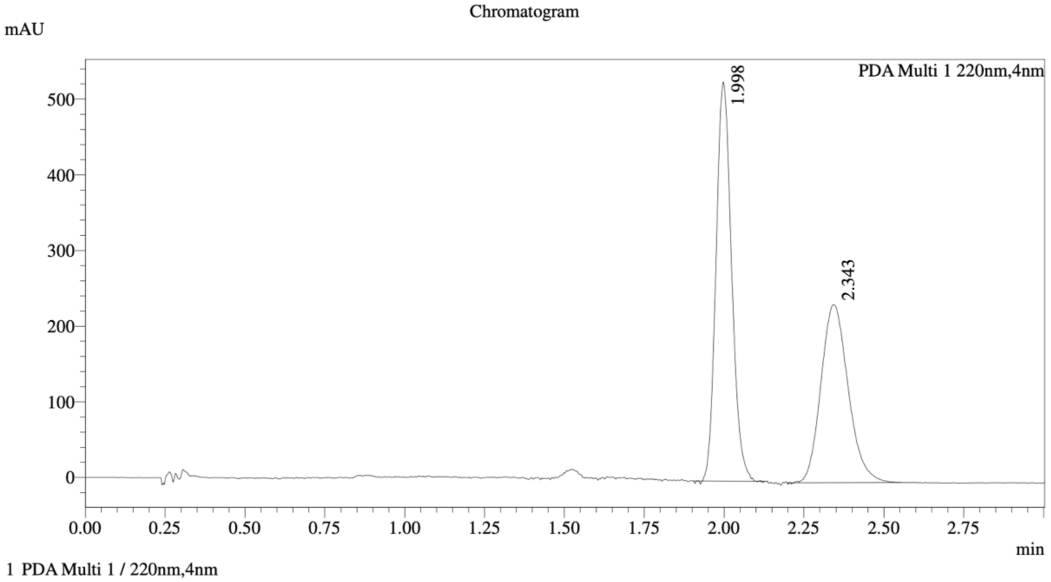


==================================================================================
Integration Result
==================================================================================

Peak Table

PDA Ch1 220nm

| Peak# | Ret. Time | Height | Height% | Resolution(USP) | Area | Area% |
|---|---|---|---|---|---|---|
| 1 | 1.998 | 524064 | 69.024 | -- | 1785090 | 55.985 |
| 2 | 2.343 | 235189 | 30.976 | 2.776 | 1403426 | 44.015 |

Method Details

Column: Chiralcel OJ-3 50×4.6mm I.D., 3 µm
Mobile phase: A: $CO_2$, B: MeOH (0.05% diethylamine)
Elution: 5% to 40% B (gradient)

WX-02-570

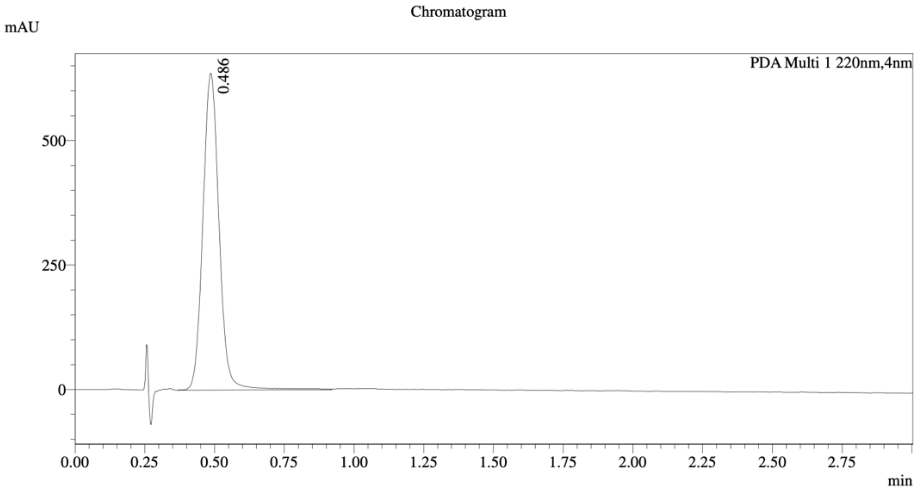


1 PDA Multi 1 / 220nm,4nm

Integration Result

Peak Table

PDA Ch1 220nm

| Peak# | Ret. Time | Height | Height% | Resolution(USP) | Area | Area% |
|---|---|---|---|---|---|---|
| 1 | 0.486 | 634232 | 100.000 | -- | 2573390 | 100.000 |

WX-02-571

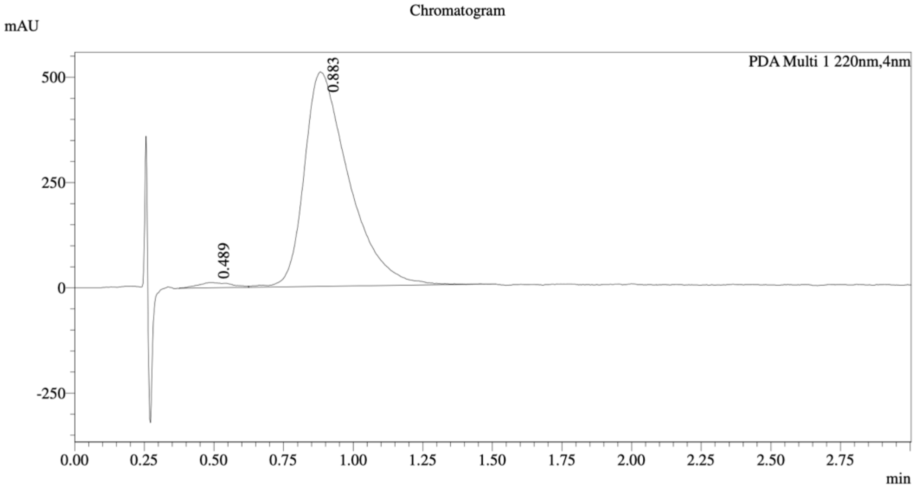


1 PDA Multi 1 / 220nm,4nm

Integration Result

Peak Table

PDA Ch1 220nm

| Peak# | Ret. Time | Height | Height% | Resolution(USP) | Area | Area% |
|---|---|---|---|---|---|---|
| 1 | 0.489 | 12459 | 2.393 | -- | 102502 | 1.771 |
| 2 | 0.883 | 508169 | 97.607 | 1.587 | 5686077 | 98.229 |

Mixture of WX-02-570 and WX-02-571

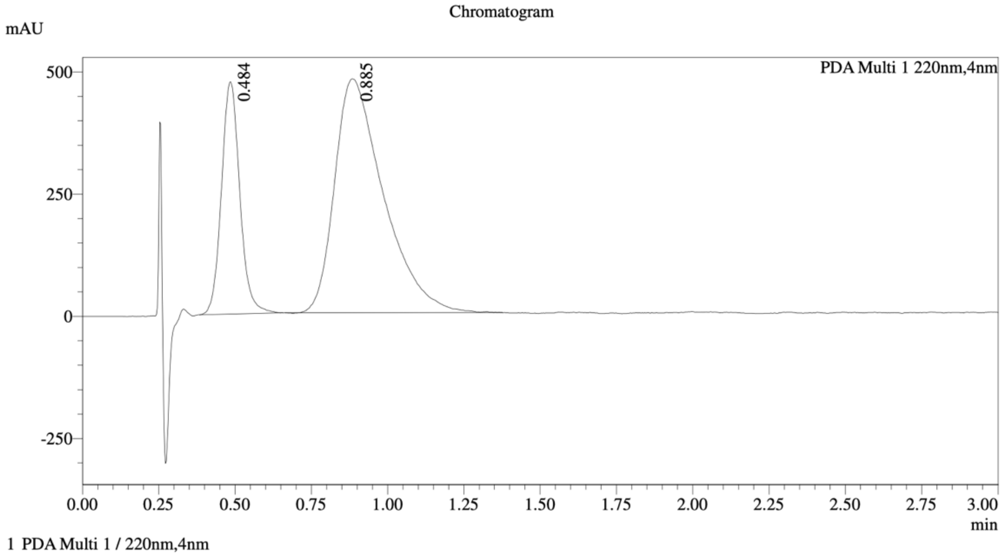


==========================================================================================

Integration Result

==========================================================================================

Peak Table

PDA Ch1 220nm

| Peak# | Ret. Time | Height | Height% | Resolution(USP) | Area | Area% |
|---|---|---|---|---|---|---|
| 1 | 0.484 | 473932 | 49.775 | -- | 1987842 | 27.133 |
| 2 | 0.885 | 478224 | 50.225 | 2.003 | 5338577 | 72.867 |

Method Details

Column: Chiralcel OD-3 50×4.6mm I.D., 3 µm
Mobile phase: A: $CO_2$, B: 2:1 MeOH/acetonitrile (0.05% diethylamine)
Elution: 40% B
